# A microglia-CD4^+^ T cell partnership generates protective anti-tumor immunity to glioblastoma

**DOI:** 10.1101/2022.08.12.502093

**Authors:** Dan Chen, Siva Karthik Varanasi, Toshiro Hara, Kacie Traina, Bryan McDonald, Yagmur Farsakoglu, Josh Clanton, Shihao Xu, Thomas H. Mann, Victor Du, H. Kay Chung, Ziyan Xu, Victoria Tripple, Eduardo Casillas, Shixin Ma, Carolyn O’Connor, Qiyuan Yang, Ye Zheng, Tony Hunter, Greg Lemke, Susan M. Kaech

## Abstract

The limited efficacy of immunotherapies against glioblastoma illustrates the urgent need to better understand the interactions between the central nervous system and the immune system. Here, we showed that a protective response to αCTLA-4 therapy depended on a mutualistic relationship between microglia and CD4^+^ T cells. Suppression of gliomas by CD4^+^ T cells did not require tumor-intrinsic MHC-II expression, but rather was dependent on the selective expression of MHC-II and antigen presentation by local microglia that in turn, sustained CD4^+^ T cell tumoricidal effector functions. CD4^+^ T cell secretion of IFNγ made the glioma cells vulnerable to enhanced tumor surveillance and phagocytosis by microglia via the AXL/MER tyrosine kinase receptors that were necessary for tumor suppression. This work illustrates a novel partnership between CD4^+^ T cells and microglia that unleashes the tumoricidal properties of microglia that can be harnessed to improve immunotherapies for glioblastoma.

## Introduction

Glioblastoma is the most common and deadliest form of brain cancer with no effective therapy available (Tan et al., 2020; Taylor et al., 2019). Glioblastoma is composed of heterogeneous populations of glioma stem cells (GSCs) that demonstrate strong resistance to most conventional and targeted therapies (Qazi et al., 2017). Single-cell profiles of human glioblastoma identified four dominant subpopulations of cancer cells including neural progenitor-like (NPC-like), oligodendrocyte progenitor-like (OPC-like), astrocyte-like (AC-like), and mesenchymal-like (MES-like), each of which is associated with well-established GSC markers such as CD24, PDGFRA, EGFR, and CD44, respectively (Hara et al., 2021; Neftel et al., 2019). Given these complexities and therapy-resistant features of GSCs, identifying more effective therapies to target heterogeneous glioblastoma GSCs is an unmet clinical need. Immunotherapy, which aims to harness the immune system’s anti-tumor capabilities, has shown impressive results in the treatment of various tumor types like melanoma, and lung cancer (Bonaventura et al., 2019; Farkona et al., 2016; Hellmann et al., 2018; Sade-Feldman et al., 2018; Zhang and Zhang, 2020), which inspired multiple clinical trials to treat late-stage glioblastoma using immune checkpoint blockade (ICB) with αCTLA-4 and/or αPD-1 (Curry and Lim, 2015; Huang et al., 2017; Lim et al., 2018; Muftuoglu and Liau, 2020). However, little clinical benefit was observed (Jackson et al., 2019; Omuro et al., 2018; Reardon et al., 2020). Thus, the present failures of ICB in glioblastoma has left the field void of a fundamental understanding of what types of immune responses and which types of immune cells will provide protection against glioblastoma and could underlie ICB-mediated tumor regression. We sought to address this need by examining more physiological preclinical glioblastoma models that resemble human GSCs to ascertain if they respond to current immunotherapies, and if so, how a protective immune response could be orchestrated in the brain.

Common features of glioblastoma tumor microenvironments (TMEs) are that they are often devoid of T cells and considered immunologically “cold” (Frederico et al., 2021; Jackson et al., 2019). However, recent work has shown that MES-like glioblastoma tends to have a greater T cell infiltration (Hara et al., 2021), pointing to the possibility that some types of GSCs may be more sensitive to immunotherapy than others. There are a relatively limited number of mouse syngeneic glioma models, such as GL261 and CT2A, which are created by chemical carcinogens and do not generate identifiable GSCs. These models have a high tumor mutation burden (TMB) and respond to ICB, unlike human gliomas (Oh et al., 2014). However, glioblastoma 005 GSC-derived brain tumors used in our study have characteristics more similar to human glioblastoma, including heterogeneous GSC-like properties, invasiveness (Marumoto et al., 2009), and more closely resemble the immune-phenotypic signature of glioblastoma patients than other models (CT2A, GL261, Mut3) (Khalsa et al., 2020). Using the 005 GSC model, we sought to determine if immune responses can be generated against GSCs by ICB, and what the immune correlates of such protection may be to better inform human glioblastoma immunotherapy trials.

Eliciting a protective anti-tumor T cell response during ICB depends on interactions with antigen-presenting cells (APCs) (Bandola-Simon and Roche, 2019; Sabado et al., 2017), such as dendritic cells (DCs), to prime and activate antigen-specific CD8^+^ or CD4^+^ T cells (Cohen et al., 2022; Schenkel et al., 2021), but often the APCs induce tolerance or suppress the anti-tumor T cells in cancer (Nurieva et al., 2013). The brain parenchyma does not typically harbor DCs in the resting state, yet a higher percentage of DCs in a glioma favors more protective immune responses (Datsi and Sorg, 2021; Liau et al., 2018; Srivastava et al., 2019), which indicates the importance of APCs to successful immunotherapy. Microglia, the brain resident macrophages, typically regulate neuronal activity, synaptic plasticity and memory at homeostasis, but they also serve critical roles in host defense against infections in the central nervous system (CNS) acting as APCs to T cells, engulfing microbes and infected cells (Byram et al., 2004; Goddery et al., 2021; Moseman et al., 2020; Schetters et al., 2017; Shaked et al., 2004). In contrast, microglia can also play immunosuppressive roles and promote glioma progression (Geribaldi-Doldan et al., 2020; Poon et al., 2017; Wei et al., 2013; Zhai et al., 2011). Immunosuppressive microglia, termed glioblastoma-associated microglia (GAMs), have become therapeutic targets for glioblastoma (Poon et al., 2017) with the intent to repolarize GAMs towards pro-inflammatory states to stimulate anti-tumor responses, but doing so requires a deeper mechanistic understanding of what signals and cell types cause microglia to switch between immuno- supportive and -suppressive states in glioblastoma.

Our studies show for the first time that brain-infiltrating CD4^+^ T cells are pivotal regulators of tumor clearance and anti-tumor function by microglia in glioblastoma. We found that αCTLA-4, but not αPD-1, prolonged survival of glioma-bearing mice in a manner dependent on CD4^+^ T cells, but not CD8^+^ T cells. CD4^+^ T cells exhibited expanded TCR clones with increased IFNγ production after αCTLA-4 treatment. Importantly, IFNγ^+^ CD4^+^ T cells also induced MHC-II on microglia, which served as requisite APCs to sustain anti-tumor functions of CD4^+^ T cells during ICB. Simultaneously, IFNγ^+^ CD4^+^ T cells also promoted microglial activation and tumor-surveillance that led to phagocytosing and eradication of tumor cells in an AXL/MER tyrosine kinase receptor-dependent manner. Collectively, our data reveal that the partnership between microglia and CD4^+^ T cells is a key driver for glioblastoma control, offering new therapeutic avenues for treating such a formidable disease.

## Results

### Anti-CTLA-4 treatment suppressed tumor growth and increased the ratio of CD4^+^ T helper cells in glioblastoma

To ascertain if anti-tumor immune responses against GSCs could be elicited by ICB we established gliomas in mice using 005 cells that generate tumors containing OPC- and MES-like stem cells **(Figure S1A)** and administered anti-PD-1 (αPD-1) and anti-CTLA-4 (αCTLA-4) mAbs for three weeks. As seen in human clinical trials, αPD-1 mAbs had no effect on median survival in 005 tumors, however, αCTLA-4 treatment significantly (p ≤ 0.001) extended survival by 2-3 folds **(Figure 1A)** and reduced tumor burden in most animals **(Figures S1B-C)**. We also examined the effects of ICB on the commonly used glioma model GL261 that was created by chemical carcinogenesis (Oh et al., 2014) and again greater therapeutic effects were observed by αCTLA-4 treatment as opposed to αPD-1 therapy **(Figure 1B).** Together, these results suggested the possibility that while glioblastoma may be resistant to αPD-1 therapy, it may be more intrinsically responsive to αCTLA-4; this provided an opportunity to explore the underlying immune correlates of effective immunotherapies against GSCs.

**Figure 1.**
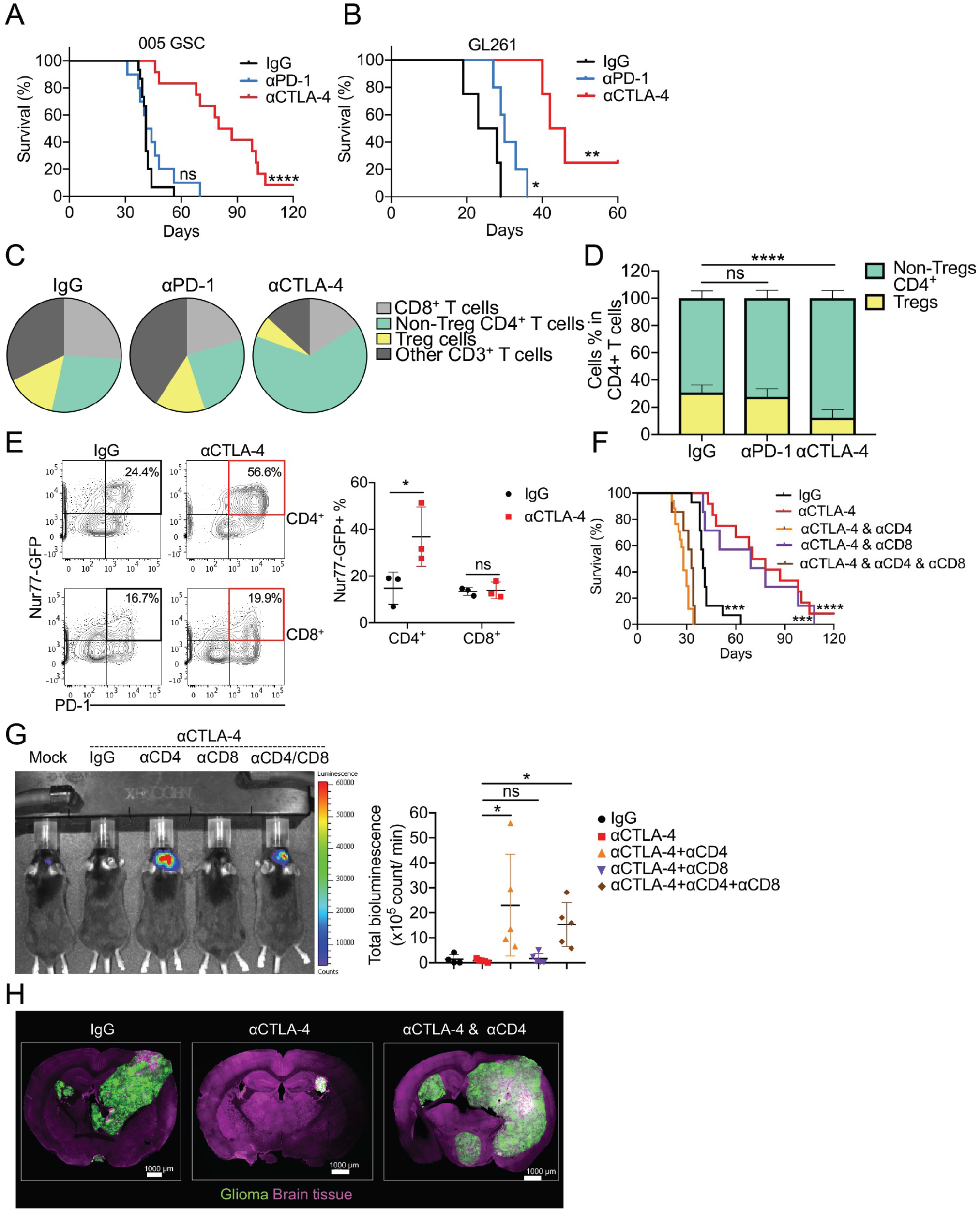
αCTLA-4 therapy prolongs glioblastoma survival in a CD4^+^ T-cell dependent manner. (A-B) C57BL/6 mice were implanted with 3 × 10^5^ 005 GSCs (A) or implanted with 5 × 10^4^ GL261 (B) on day 0, and treated with αPD-1 antibody (10 mg/kg), αCTLA-4 (10 mg/kg) or isotype control IgG (10 mg/kg) injected i.p. on days 7, 14, 21 and 28. (A) Median survival of IgG (41 days; n = 15) was compared with αCTLA-4 (83.5 days; n = 12, p = 0.000002), αPD-1 (42.5 days; n = 10, p = 0.2632) by log-rank analysis. (B) Median survival of IgG (25.5 days; n = 5) was compared with αCTLA4 (44 days; n = 6, p = 0.0067), αPD-1 (30 days; n = 5, p = 0.044) by log-rank analysis. (C) Pie charts showed the relative proportion of T cell subpopulation (CD3^+^) changes from various treated glioma samples (pie chart: CD8^+^ T cells [CD3^+^CD8^+^], Non-Treg CD4^+^ T cells [CD3^+^CD4^+^FOXP3^-^], Treg cells [CD3^+^CD4^+^FOXP3^+^], Others [CD3^+^CD4^-^CD8^-^]). (D) Frequency of Treg (% of FOXP3^+^ cells in CD4^+^ T cells) and non-Treg CD4^+^ T cells (% of FOXP3^-^ cells in CD4^+^ T cells) from glioma tissue was quantified. (E) Representative flow plots and summarized frequencies of Nur77-GFP expression in CD4^+^ or CD8^+^ T cells from mice glioma tissue. (F) Mice were implanted with 3 × 10^5^ 005 GSCs on day 0, and treated with 10 mg/kg IgG or αCTLA-4 with or without αCD4 antibody, or αCD8 antibody injected i.p. on days 7, 14, 21 and 28. Median survival of IgG (40 days; n = 14) was compared with αCTLA-4 (74 days; n = 11, p = 0.000008). Similarly, αCTLA-4 was compared with αCTLA-4 combined with αCD4 (30 days; n = 13, p < 0.000001), αCTLA-4 combined with αCD8 (69 days; n = 7, p = 0.6107), and αCTLA4 combined with αCD8 and αCD4 (33 days; n = 7, p=0.000002) by log-rank analysis. (G) Mice were implanted with 3 × 10^5^ 005-Luc GSCs on day 0, and were treated as in (F). Mice were imaged on day 28. Bioluminescence was quantified. n = 5 mice / group. (H) Mice were implanted with 3 × 10^5^ 005 GSCs, and the localization of 005 cells in the brain was analyzed by immunofluorescence confocal imaging. All scale bars indicate 1000 μm. All results were pooled from or representative of 2-4 experiments with n = 4 mice / group (C), n = 5-13 mice / group (D), n = 3 mice / group (E), n = 5 mice / group (G), n = 4 mice / group (H). Data are expressed as mean ± SEM. Statistical analysis was performed using Student’s *t* test (two tailed) comparing IgG group to various treatment groups. * p < 0.05, ** p < 0.01, *** p < 0.001, **** p < 0.0001.

To examine the immune cell types that correlate with ICB-response or resistance, we used flow cytometry to profile numerous immune cell populations in 005 tumors after αPD-1or αCTLA-4 treatment. αPD-1 had minimal effects on the composition and frequencies of T cells compared to the IgG control correlating with the reduced efficacy **(Figures 1C and S1D)**. In contrast, αCTLA-4 treatment substantially increased the frequency of tumor-infiltrating CD4^+^ T helper cells and reduced the frequency of Tregs **(Figures 1C-D, and S1D-E)**. Using the Nur77^GFP^ reporter mouse (Moran et al., 2011), a transcriptional reporter for T cell receptor activation, we observed a profound increase in the percentage of Nur77-GFP^+^ CD4^+^ T cells relative to CD8^+^ T cells after αCTLA-4 treatment **(Figure 1E)**, demonstrating the more selective influence of αCTLA-4 treatment on tumor-reactive CD4^+^ T helper cells as seen in other tumor models (Wei et al., 2017). Together, these results show that αCTLA-4 treatment greatly improved survival, suppressed tumor growth, and increased the frequency and activation of CD4^+^ T helper cells in tumor-bearing mice.

To examine if the therapeutic effects of αCTLA-4 ICB were dependent on T cells, we depleted CD4^+^ and CD8^+^ T cells, and this showed that depletion of CD4^+^ T cells abrogated the therapeutic effects of αCTLA-4 therapy, but depletion of CD8^+^ T cells did not **(Figures 1F and S1F)**. The requirement of CD4^+^ T cells in glioma regression following αCTLA-4 treatment was also evident by imaging of luciferase^+^ or GFP^+^ 005 using IVIS **(Figure 1G)** or confocal microscopy (**Figure 1H**), respectively. Together, these results indicated that harnessing CD4^+^ T cell function can be a highly protective and critical form of anti-tumor immunity in preclinical models of glioma that better resembles the heterogeneity of human glioblastoma.

### Th1-like CD4^+^ T helper cells were essential for anti-tumor immunity and longer-term survival in glioblastoma

Since CD4^+^ T cells played a critical role in anti-tumor immunity in glioma-bearing mice, we next profiled intratumoral T cell repertoire after αCTLA-4 treatment using paired single-cell (sc) T cell receptor (TCR)- and RNA-sequencing. A total of 6,174 T lymphocytes were identified with matched gene expression and V(D)J combination profiles and visualized using Uniform Manifold Approximation and Projection (UMAP). Three primary T cell clusters were distinguished: (1) CD8^+^ T cells (*Cd8a*^+^), (2) Treg (*Cd4*^+^ *Foxp3*^+^), and (3) Non-Treg CD4^+^ T cells (*Cd4*+*Foxp3*^-^) **(Figures 2A and S2A)**. Examination of the scTCR-seq data showed an array of expanded clonotypes in mainly CD8^+^ and Non-Treg CD4^+^ T cell subsets after αCTLA-4 treatment **(Figure 2B)**. The majority of the CD4^+^ T cell TCR clonotypes expanded by αCTLA-4 treatment were novel compared to the IgG controls **(Figure 2C)**. However, one TCR clonotype [CAASGGSNYKLTF_CASSSGEDTQYF] present in IgG group expanded to become the predominant clonotype in αCTLA-4 treated tumors **(Figure 2C)** and this clonotype has also been reported in GL261 glioma model (Mohme et al., 2020), highlighting a possible shared MHC-II- restricted glioma-associated antigen. Moreover, examination of the top 10 most abundant clones after αCTLA-4 treatment showed predominant *Ifng* and T-bet *(Tbx21)* expression within these cells, indicating expansion of tumor specific effector T helper 1 (Th1) cells after ICB **(Figure 2D)**. Accordingly, paired transcriptome analysis demonstrated that several hallmark Th1 genes expressed in CD4^+^ tumor-infiltrating lymphocytes (TILs) were induced by αCTLA-4 therapy relative to IgG controls including cytokine and chemokine genes (*Ifng, Il2, Il21, Csf2, Cxcl13*), chemokine receptors *(Cxcr3, Cxcr6, Ccr8),* co-stimulatory genes (*Cd28, Cd40lg*), transcription factors (*Tbx21, Bhlhe40, Nr4a1, Id2*) **(Figure 2E)**. While CD8^+^ T cells and Tregs from αCTLA- 4 treated gliomas also displayed unique TCR repertoires compared to IgG control-treated tumors **(Figure S2B)**, fewer significant changes in gene expression were observed in CD8^+^ TILs and Tregs.

**Figure 2.**
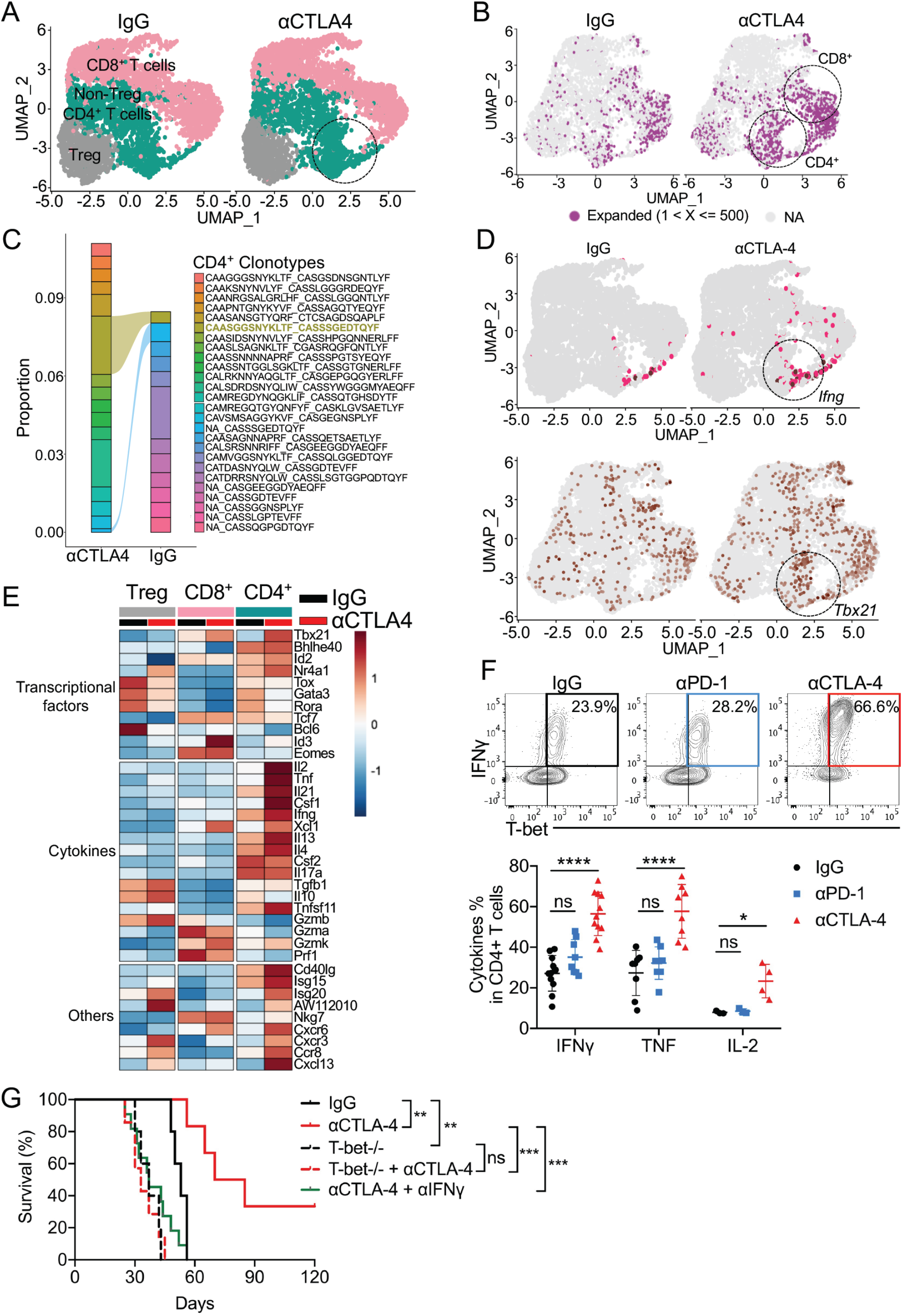
Th1 cells are essential for anti-tumor immunity and long-term survival in glioblastoma. (A) scRNAseq and scTCRseq were performed in brain harvested T cells from 005 bearing mice treated with IgG or αCTLA-4 (35 days post implantation). The total number of recovered cells was 8,139 cells from scRNA-seq, and 5,514 T lymphocytes from V(D)J combination information for αCTLA-4 treated group, and 9,746 cells from scRNA-seq data and 3,279 T lymphocytes from V(D)J combination information for IgG treated group. Uniform manifold approximation and projection (UMAP) displayed 3 distinct clusters, including the clustering of CD8^+^ T cells (CD3^+^CD8^+^), Treg (FOXP3^+^CD4^+^) and non-treg CD4+ T cells (FOXP3^-^CD4^+^) by integrated analysis. (B) The highly expanded clonotypes was displayed overlaid with the UMAP in each subset from A. (C) Shared clonotype analysis in non-Treg CD4^+^ T cells between IgG vs. αCTLA-4 treatment by alluvial clonotypes. (D) Feature plot showed *Ifng* and *Tbx21* in the different clusters between IgG vs. αCTLA-4 as defined in A. (E) Heatmap showed the expression dynamics of transcription factors, cytokines and others enriched in the different clusters between IgG vs. αCTLA-4. (F) C57BL/6 mice were implanted with 3 × 10^5^ 005 GSCs on day 0, were treated as in (Figure 1A), and euthanized on day 35. The expression of IFNγ, TNF, IL-2 was measured and quantified by flow cytometry in CD4^+^ T cells from harvested brains. (G) T-bet^+/+^ or T-bet^-/-^ mice were implanted with 3 × 10^5^ 005 GSCs on day 0, treated with 10 mg/kg IgG or αCTLA-4 combined with or without αIFNγ i.p. on days 7, 14, 21 and 28, and monitored for survival. Median survival of IgG (53 days; n = 5) was compared with αCTLA-4 (77.5 days; n = 6, p = 0.00369). Median survival of T-bet^-/-^ (37 days, n=5) was compared with T-bet^-/-^ treated with αCTLA-4 (33 days; n = 7, p = 0.85176). Similarly, comparison of αCTLA-4 treatment of wild-type B6 mice with that of T-bet^-/-^ (33 days; n = 7, p = 0.00036), as well as the corresponding comparison in αCTLA-4 combined with αIFNγ (43 days; n = 9, p = 0.00045), were done by log-rank analysis. All the results were pooled from or representative of 2-4 experiments with n = 3-5 mice / group (A-E), n = 4-11 mice / group (F). Data are expressed as mean ± SEM. Statistical analysis was performed using Student’s *t* test (two tailed) comparing IgG group to various treatment groups. * p < 0.05, ** p < 0.01, *** p < 0.001, **** p < 0.0001.

In agreement with the mRNA expression, αCTLA-4 treatment, but not αPD-1 treatment, significantly increased the proportion of PD-1^+^ T-bet^+^ CD4^+^ TILs that produce interferon-γ (IFNγ), tumor necrosis factor (TNF) and interleukin-2 (IL-2) (**Figure 2F**). Further, treatment with αCTLA-4 substantially increased the amount of IFNγ and TNF produced on a per cell basis (mean fluorescent intensity (MFI)) within CD4^+^ T cells (**Figure S2C**). The CD8^+^ TILs also showed increased TNF and only modestly increased granzyme B (GzmB) and IFNγ with αCTLA-4, despite not being required for the therapeutic effects **(Figure S2D)**. To test if T-bet or IFNγ were required for the protective effects of anti-glioblastoma CD4^+^ T cells, we implanted 005 GSC tumors in T-bet knockout mice (*Tbx21^-/-^*) and treated them with or without αCTLA-4. This showed the therapeutic benefit of αCTLA-4 was lost in T-bet-deficient mice **(Figure 2G**) and correlated with a lack of IFNγ^+^ CD4^+^ T cells in the tumors **(Figure S2E)**. Similarly, IFNγ blockade abrogated the response to αCTLA-4 **(Figure 2G)**. Collectively, these results show that the immune protection provided by αCTLA-4 treatment in glioblastoma is largely based on a dominant and robust IFNγ-producing Th1 CD4^+^ T cell response consisting of a largely novel TCR repertoire.

### Microglia promote anti-glioblastoma CD4^+^ T cell responses and repress glioma growth

In order to generate and sustain a protective CD4^+^ T cell response to ICB, we next asked which cell types express MHC-II in the brain. Examination of the scRNA-seq identified infiltrating DCs, B-cells, macrophages and brain-resident microglia as the four primary MHC-II expressing cell populations out of 11 clusters **(Figures 3A-B, S3A)**, which was also confirmed by flow cytometry **(Figure S3B)**. To interrogate which of these populations supported a protective anti-tumor CD4^+^ T cell response, we systematically examined the role of each MHC- II expressing population. First, to separate the role of conventional DCs (cDCs) from initial priming vs. sustaining CD4^+^ T cell responses in the CNS, we implanted tumors in CD11c-DTR mice and *14 days later*, began diphtheria toxin (DT) treatment to deplete CD11c^+^ cells **(Figure 3C)**. Surprisingly, depletion of CD11c^+^ cDCs did not influence the infiltration or function of CD4^+^ T cells **(Figures 3C-E and S3C)**. Similarly, depletion of CD169^+^ macrophages using CD169-DTR mice also had minimal effect on the density and function of CD4^+^ T cell in CNS **(Figures S3D-E)**. Additionally, αCTLA-4 treatment to *mu*MT^-^ (B cell deficient) **(Figures S3F- G)** or *Ccr2*^-/-^ (defective monocyte recruitment) **(Figures S3H-I)** mice had minimal impact on efficacy of αCTLA-4 and IFNγ production of CD4^+^ TIL **(Figures S3G and S3I)**, albeit the *Ccr2^-/-^* mice showed some reduction in efficacy compared to controls **(Figure S3H)**. Together, these data suggest that neither B cells nor bone marrow-derived monocytes/macrophages nor conventional DCs were necessary for sustaining an anti-glioblastoma CD4^+^ TIL response.

**Figure 3.**
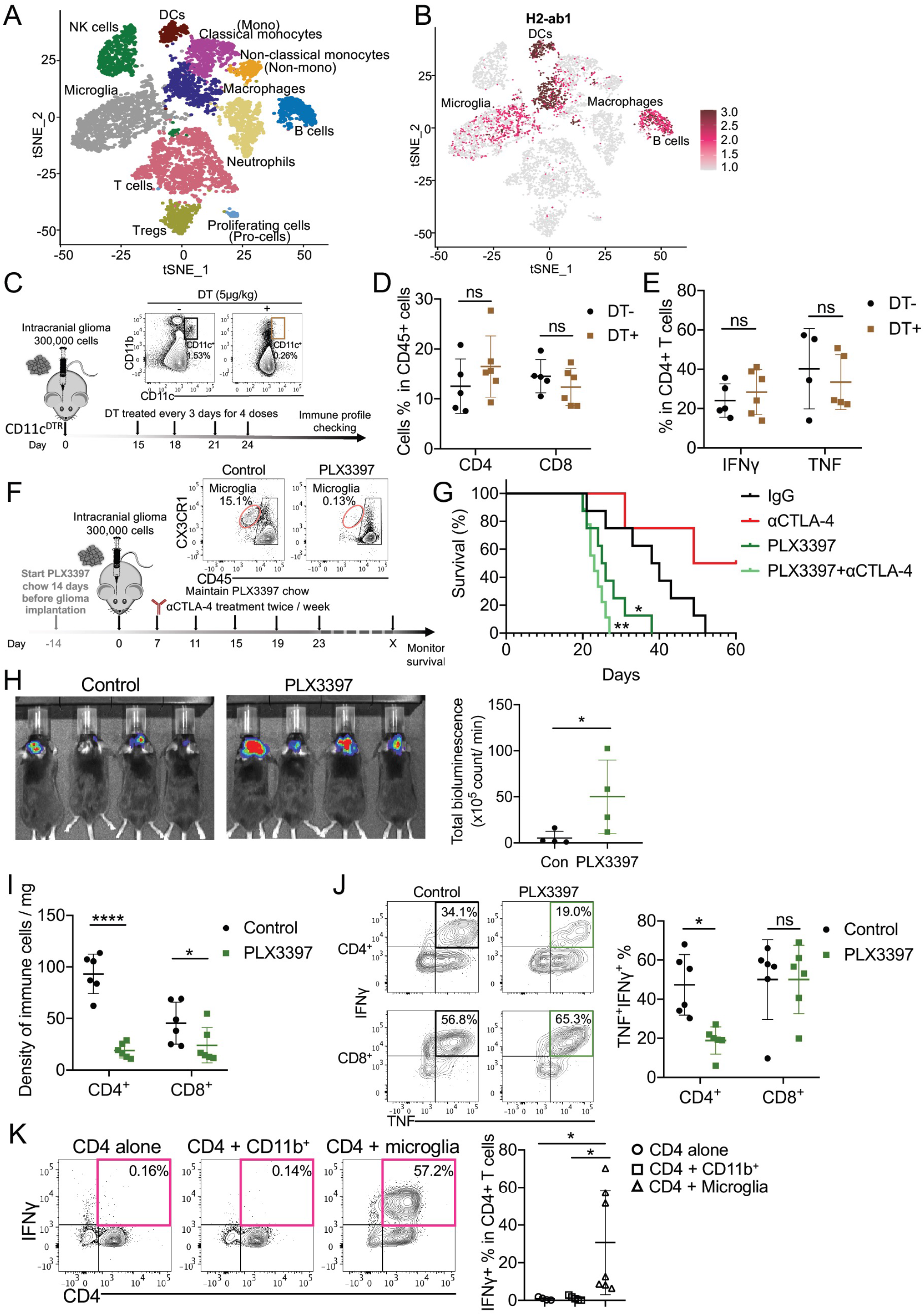
Microglia, but not CD11c^+^ cells, are required for anti-glioblastoma immune response. (A) tSNE plot of CD45^+^ clusters from merged conditions. Microglia (CD45^int^CX3CR1^+^), myeloid cells (CD45^hi^CD11b^+^), T cells (CD45^hi^CD3^+^) and all remaining immune cells (CD45^hi^CD11b^-^CD3^-^) were sorted from normal brain and 005 glioma-bearing brains treated with IgG, anti-CTLA-4, or αCTLA4 combined with CD4 depletion. Cells from each treatment group were pooled at equal ratios and then subjected to scRNA-seq. We recovered 1,420 immune cells from normal brain, 1,484 immune cells from IgG group, 1,874 immune cells from αCTLA4 groups, and 3,916 immune cells from αCTLA4 combined with CD4 depletion groups. Eleven clusters were identified on the basis of previously described cell-type specific gene expression patterns. (B) Feature plots of *H2-ab1* gene. (C-E) CD11c-DTR mice were administered with 5 μg/kg DT (i.p.) 2 weeks after glioma implantation, and maintained the treatment twice/week for 2 weeks. The efficacy of depletion of CD11c^+^ cells was checked by flow cytometry (C). The percentages of CD4^+^ and CD8^+^ T cells (D), and PD-1^+^IFNγ^+^ and PD-1^+^TNFα^+^ of CD4^+^ cells (E), in brain tumor tissues from different treatment groups were measured and quantified by flow cytometry. (F-H) C57BL/6 mice received control chow or PLX3397 chow (290 mg/kg) for 14 days prior to implantation, and then were implanted with 3 × 10^5^ 005 GSCs (F-G), or 005-Luc GSCs (H). The depletion efficiency of microglia (CD45^int^CX3CR1^+^) was assessed by flow cytometry (F). Median survival of IgG (39 days; n = 8) was compared with PLX3397 (25.5 days; n = 8, p = 0.01135). Similarly, αCTLA-4 (54.5 days; n = 4) was compared with αCTLA-4 combined with PLX3397 (23 days; n = 9, p = 0.00197) by log-rank analysis (G). Bioluminescence was quantified (H). (I) The densities of CD8^+^ and CD4^+^ T cells in control or PLX3397 treated groups from 005 tumors were quantified by flow cytometry. (J) The expression of IFNγ^+^TNF^+^ was measured and quantified in CD4^+^ and CD8^+^ T cells in control or PLX3397 treated groups from 005 tumors. (K) CD4^+^ T cells (CD45^+^CD4^+^), microglia (CD45^int^CX3CR1^+^), and CD11b^+^ myeloid (CD45^hi^CD11b^+^) cells were isolated from glioma-bearing brains and co-cultured *in-vitro* in the presence of GolgiStop for 6 hours as indicated. IFNγ production was measured by flow cytometry. All results were pooled from or representative of 2-4 experiments with n = 3-5 mice / group (A-B), n = 5 mice / group (C-E), n = 4 mice / group (H), n = 6 mice / group (I-J), n = 5-7 mice / group (K). Data are expressed as mean ± SEM. Statistical analysis was performed using Student’s *t* test (two tailed) comparing IgG group to various treatment groups. * p < 0.05, ** p < 0.01, *** p < 0.001, **** p < 0.0001.

Lastly, we examined the role of microglia in supporting anti-tumor CD4^+^ TILs by depleting microglia using a CSF1R inhibitor (PLX3397). Mice were fed with PLX3397 chow for 14 days until tumor implantation, which achieved complete depletion of microglia **(Figure 3F)**, but did not significantly influence other MHC-II^+^ myeloid populations **(Figures S3J-K)**. Importantly, depletion of microglia significantly reduced survival **(Figure 3G)** and enhanced glioma growth **(Figure 3H)** in mice treated with or without αCTLA-4. Additionally, loss of microglia by PLX3397 dramatically impaired the expansion of IFNγ^+^ TNF^+^ CD4^+^ T cells whereas the effects on CD8^+^ T cells were rather modest **(Figures 3I-J)**. To independently compare the ability of microglia or other local myeloid cells to reactivate tumor-infiltrating CD4^+^ T cells *in vitro*, we sorted microglia and CD11b^+^ myeloid cells from the brains of tumor bearing mice and incubated them for twelve hours with CD4^+^ T cells sorted from the same tumor bearing mice. Microglia induced the strongest reactivation of CD4^+^ T cells based on IFNγ expression as measured by flow cytometry whereas the other local myeloid populations failed to do so **(Figure 3K)**. These results suggest that microglia are important for sustaining the function of anti-glioblastoma CD4^+^ T cells upon αCTLA-4 treatment in the brain.

### Microglia directly interact with CD4^+^ TILs via MHC-II to sustain their anti-glioblastoma effector responses during ICB

Since microglial depletion was associated with reduced CD4^+^ T cells density and function **(Figures 3I-J)**, we hypothesized that microglia play a direct role in sustaining activated CD4^+^ TILs and effector functions locally in the brain via tumor antigen-presentation. We observed that microglia upregulate MHC-II and IL-12 in response to αCTLA-4 therapy **(Figures 4A-B and S4A**), suggesting microglia provide support to Th1 effector functions in the brain. To explore whether microglial MHC-II was required to sustain anti-GMB CD4^+^ TILs, we crossed *Tmem119 Cre-ERT2* mice (Kaiser and Feng, 2019) to those carrying a floxed *H2-ab1* allele and then treated *H2-ab1^fl/fl^ Tmem119^CreERT2/+^* mice with tamoxifen one week before glioma implantation to delete MHC-II selectively in CD45^int^ CX3CR1^+^microglia **(Figures 4C** and **S4B)**. We then implanted 005 tumors into *H2-ab1^fl/fl^ Tmem119^CreERT2/+^* mice and littermate controls (*H2-ab1^+/+^ Tmem119^CreERT2/+^*) and treated with αCTLA-4 and tamoxifen for an additional 2 weeks (**Figure 4C**). We validated that MHC-II expression was absent from CD45^int^ microglia, but was expressed by the other bone-marrow derived APCs that infiltrate the brain (**Figures 4C and S4B-C**). Importantly, this showed that selective depletion of MHC-II in microglia ablated the therapeutic response to αCTLA-4 therapy as assessed by survival **(Figure 4D)** and tumor growth **(Figure S4D)**. The most prominent effects of deleting MHC-II from microglia was a significant reduction in the density of IFNγ^+^PD-1^+^ CD4^+^ T cells **(Figures 4E-F)**. Similar results were also observed using an alternative approach of generating brain-protected irradiated bone marrow chimeras (BMCs) by transferring WT bone marrow (BM) into MHC-II KO mice or WT mice **(Figures 4G-H** and **S4E-F)**. After four months of bone marrow reconstitution, MHC-II KO animals that received WT BM had MHC-II^-/-^ microglia, but MHC-II^+^ peripheral antigen-presenting cells (APCs). Conversely, WT animals that received WT BM had normal MHC-II^+^ microglia, as well as MHC-II^+^ peripheral APCs **(Figure S4E)**. In the BMCs lacking MHC-II^+^ microglia (i.e., the WT➔MHC-II^-/-^ BMCs), there was a dramatic decrease in the density of CD4^+^ T cells within CNS of, whereas CD8^+^ T cell density was only slightly reduced **(Figure S4F)**. Moreover, absence of MHC-II in microglia significantly impaired the expression of T-bet and IFN-γ on CD4^+^ T cells, indicating that MHC-II^+^ microglia are important for sustaining CD4^+^ Th1 anti-tumor immunity **(Figure 4H)**. Altogether, these findings clearly demonstrate that MHC-II^+^ microglia have a superior ability to present tumor antigens to CD4^+^ T cells compared to other myeloid cells in gliomas and that local MHC-II^+^ microglia are necessary to sustain protective IFNγ-producing anti-tumor CD4^+^ TILs.

**Figure 4.**
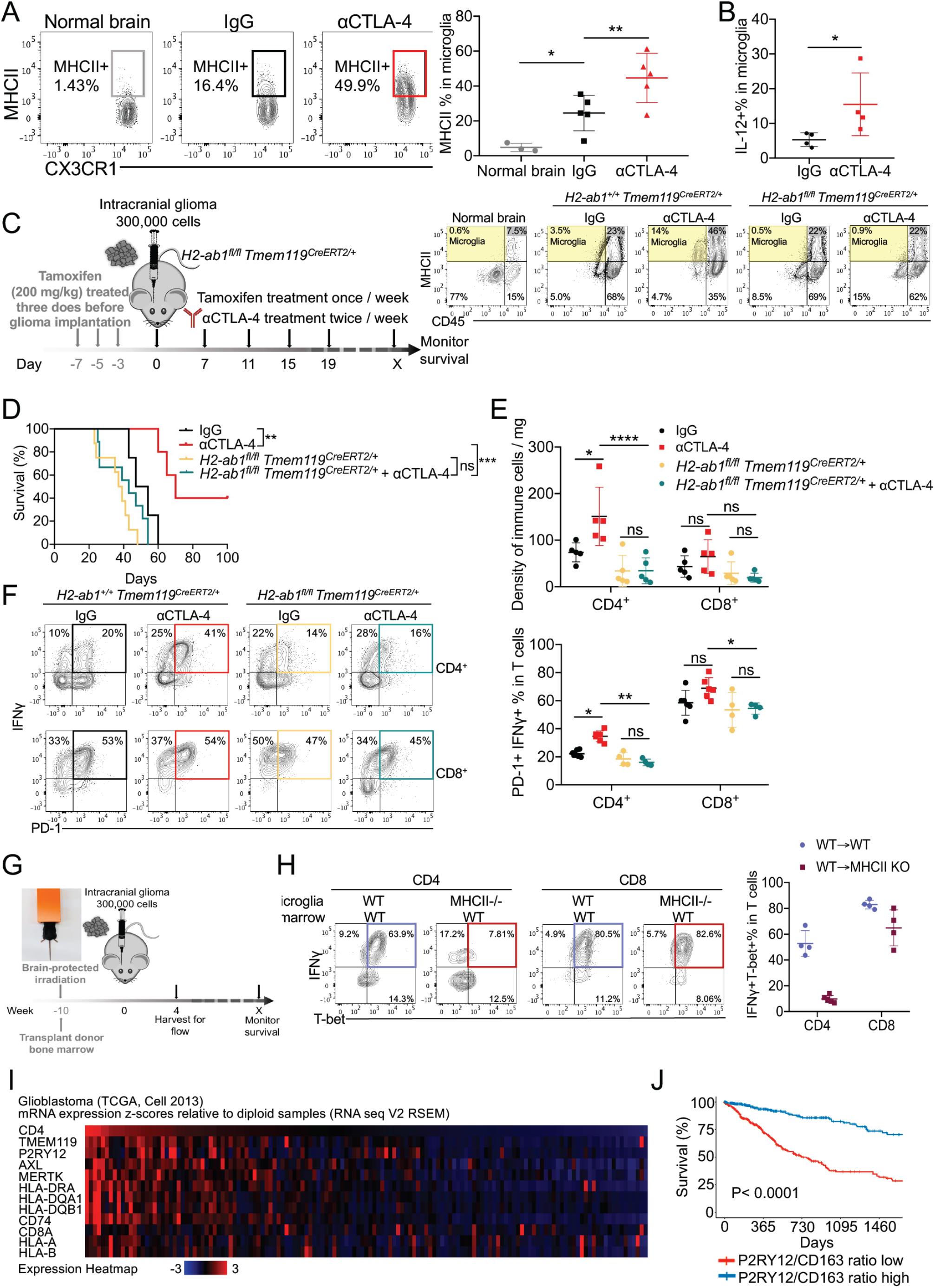
MHC-II on microglia is required for CD4^+^ T cells activation and tumor control. (A) C57BL/6 mice were implanted with PBS (normal brain without tumor as control) or 3 × 10^5^ 005 GSCs on day 0, treated with 10 mg/kg αCTLA-4 or IgG injected i.p. on days 7, 14, 21, 28, and euthanized on day 35. The expression of MHC-II was measured and quantified by flow cytometry in microglia (CD45^int^CX3CR1^+^) from harvested brains. (B) IL-12^YFP^ reporter mice were implanted with 3 × 10^5^ 005 GSCs on day 0, treated with 10 mg/kg αCTLA-4 or IgG injected i.p. on days 7, 14, 21, 28, and euthanized on day 35. The percentage of IL-12 in microglia was quantified. (C-F) *H2-ab1^fl/fl^Tmem119^CreERT2/+^* mice were treated with 200 mg/kg tamoxifen for 1 week, and then implanted with 3 × 10^5^ 005 GSCs on day 0, treated with 10 mg/kg IgG or αCTLA-4 on days 7, 14, 21 and 28, and monitored for survival. The efficacy of depletion of MHC-II on microglia was checked by flow cytometry (C). Median survival of IgG (50.5 days; n = 4) was compared with αCTLA-4 (70 days; n = 5, p = 0.00717). Median survival of *H2-ab1^fl/fl^Tmem119^CreERT2/+^*(39 days, n=7) was compared with *H2-ab1^fl/fl^Tmem119^CreERT2/+^*treated with αCTLA-4 (49 days; n = 6, p = 0.022187). Similarly, wild-type B6 mice treated with αCTLA-4 was compared with *H2-ab1^fl/fl^Tmem119^CreERT2/+^*treated with αCTLA-4 (p = 0.00191) by log-rank analysis (D). The density CD4^+^ and CD8^+^ T cells (E), and PD-1^+^ IFNγ^+^ of CD4^+^ or CD8^+^ T cells (F) in brain tumor tissues from different treatment groups were measured and quantified by flow cytometry. (G-H) Bone marrow (BM) chimeras were generated by using C57BL/6 (Ly5.1) BM injected into Ly5.1 or MHC-II KO (C57BL/6 genetic background) mice at 5 × 10^6^ cells per recipient 4 hours after lead shield brain-protected 1200 Rads irradiation (G). Brain tissues were harvested on day 35 after 005 implantation, and T-bet^+^ IFNγ^+^ of CD4^+^ or CD8^+^ T cells (H) in brain tumor tissues were measured and quantified by flow cytometry. (I) Heatmap of indicated genes in The Cancer Genome Atlas Glioblastoma Multiforme (TCGA- glioblastoma) data. (J) Survival analysis of glioma patients based on P2RY12/CD163 ratio. Overall survival (OS) is shown for P2RY12/CD163 low-ratio patients (red) against P2RY12/CD163 high-ratio patients (blue). Patient data gathered from the TCGA databases. All results were pooled from or representative of 2-4 experiments with n = 3-5 mice / group (A), n = 4 mice / group (B), n = 4-6 mice / group (E-F), n = 4 mice / group (G-H). Data are expressed as mean ± SEM. Statistical analysis was performed using Student’s *t* test (two tailed) comparing IgG group to various treatment groups. * p < 0.05, ** p < 0.01, *** p < 0.001, **** p < 0.0001.

Analysis of human glioblastoma transcriptomes deposited to the cancer genome atlas (TCGA) (Brennan et al., 2013; Ceccarelli et al., 2016) similarly showed that gliomas with abundant microglia and MHC-II expression have the greatest infiltration of CD4^+^ TILs. That is, tumors containing greater amounts of microglial activation signature genes (*P2RY12*, *TMEM119, CD74, AXL, MERTK)* were similarly enriched with mRNAs for *CD4* and MHC-II genes (*HLA-DRA, -DQA1, -DQB1)*, but not *CD8A* or MHC-I genes *(HLA-A, -B)* **(Figures 4I and S4G-I)**. Furthermore, patients whose tumors had a higher ratio of microglia-specific transcripts (*P2RY12*) to tumor associated macrophage (TAM)-specific transcripts (*CD163*) displayed longer overall survival **(Figure 4J)**, suggestive of microglial anti-tumor activities. This is in line with recent studies that analyzed distinctive microglia markers and suggested that microglia are protective against glioblastoma (Chicoine et al., 2007; Hwang et al., 2009; Sarkar et al., 2014; Sarkar et al., 2020; Woolf et al., 2021).

### IFN**γ**^+^ CD4^+^ T cells, but not CD8^+^ T cells, are necessary to increase MHC-II in glioma associated microglia

Next, we asked if the upregulation of MHC-II on microglia in gliomas depended on CD4^+^ T cells and IFNγ production. First, we compared the differentially expressed genes (DEGs) between microglia isolated from *(i)* αCTLA-4 *vs*. *(ii)* IgG *vs. (iii)* αCTLA-4 + αCD4^+^ T cell depletion treated animals using scRNA and bulk mRNA sequencing. This showed that MHC-II genes (*H2- Ab1*, *H2-Eb1, H2-Aa, Cd74*) were the highest upregulated genes in microglia after αCTLA-4 treatment (**Figure S5A**) and were highly dependent on CD4^+^ T cells (**Figure 5A and S5B**). By Metascape (Zhou et al., 2019) analysis, all DEGs from αCTLA-4 vs. αCTLA-4+αCD4 in microglia were highly enriched in several categories such as antigen processing and presentation, response to IFNγ and regulation to defense response (**Figure 5B**). The dependence on CD4^+^ T cells for microglial MHC-II upregulation was confirmed by flow cytometry (**Figure 5C**). Interestingly, induction of MHC-II by CD4^+^ T cell during ICB was fairly selective to microglia because expression of MHC-II on DCs, B cells or macrophages in the brain did not decline very much in the absence of CD4^+^ T cells (**Figures 5D-E**). This CD4^+^ T cell-microglia interaction was also computationally supported by NicheNet (Bonnardel et al., 2019; Browaeys et al., 2020) that examines gene regulatory effects between putative interacting cell populations using scRNA-sequencing data. This analysis highlighted the IFNγ signal from CD4^+^ T cells as a potential upstream signal driving microglia activation by upregulating *H2-Eb1* and *Ifitm3* levels **(Figure S5C)**.

**Figure 5.**
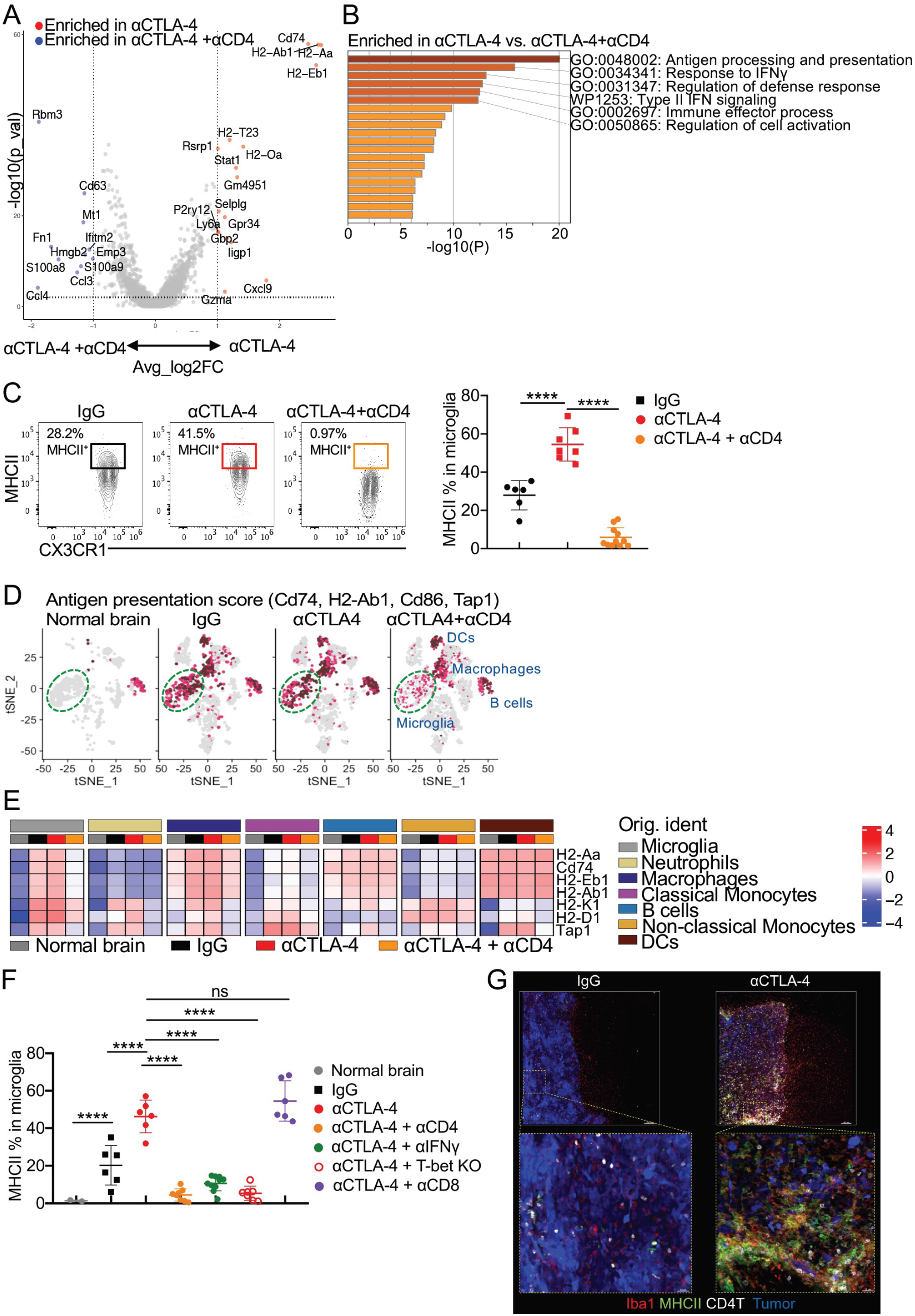
CD4^+^ T cells regulate MHC-II expression on microglia. (A) Volcano plots of genes present the magnitude (log2 (fold change), x-axis) and significance (− log10 (adjusted P value), y-axis) for microglia from αCTLA-4 vs. αCTLA-4+αCD4 from sc- RNAseq. Each spot represents a transcript. Two vertical dashed lines represent the threshold of fold changes (log2 (fold change)>1 or< −1). (B) Bar chart of clustered enrichment ontology categories was performed by Metascape on all (differentially expressed genes) DEGs in microglia from αCTLA-4 versus αCTLA-4+αCD4. (C) C57BL/6 mice were implanted with 3 × 10^5^ 005 GSCs on day 0, and treated with 10 mg/kg isotype IgG or αCTLA-4 with or without αCD4 antibody injected i.p. on days 7, 14, 21 and 28. Representative flow plots of the expression of MHC-II on microglia from tumor tissues in different groups were measured and quantified by flow cytometry. (D) tSNE plots of antigen presentation associated markers (*Cd74, H2-Ab1, Cd 86, Tap1*) from all the clusters in each condition (Normal brain, IgG, αCTLA-4, and αCTLA-4+αCD4). (E) Heatmap showed the expression dynamics of selected genes enriched in the different myeloid clusters from each condition. (F) T-bet^-/-^ mice or C57BL/6 mice were implanted with 3 × 10^5^ 005 GSCs on day 0, treated with 10 mg/kg IgG, or αCTLA-4 with or without αCD4, or αCD8 depletion antibodies, or αIFNγ blocking antibody injected i.p. on days 7, 14, 21 and 28. The MHC-II expression on microglia were measured and quantified by flow cytometry. (G) Mice were implanted with 3 × 10^5^ 005 GSCs, and the localization of Iba1^+^, CD4^+^, MCHII^+^ cells in the brain was analyzed by immunofluorescence confocal imaging. Top scale bars indicate 150 μm, and bottom scale bars indicate 30 μm. All results were pooled from or representative of 2-4 experiments with n = 3-5 mice / group (A- B, D-E), n = 6-11 mice / group (C), n = 5-8 mice / group (F), n = 4 mice / group (G). Data are expressed as mean ± SEM. Statistical analysis was performed using Student’s *t* test (two tailed) comparing IgG group to various treatment groups. * p < 0.05, ** p < 0.01, *** p < 0.001, **** p < 0.0001.

Second, we examined whether IFNγ-producing Th1 CD4^+^ T cells were important for microglia MHC-II expression using T-bet KO mice or blocking IFNγ in mice containing 005 gliomas that were treated with αCTLA-4. Both of these approaches showed that IFNγ-producing Th1 cells were specifically necessary for induction of MHC-II on microglia during ICB (**Figure 5F** and **S5D**). Interestingly, depletion of CD8^+^ T cells did not impact microglial MHC-II expression after aCTLA-4 treatment, which was surprising because the CD8^+^ T cells present in the 005 gliomas and produced IFNγ similar proportions to the CD4^+^ T cells **(Figures S2D** and **4F)**. To more directly compare the effects of CD4^+^ and CD8^+^ T cells in regulating microglial antigen presentation in gliomas we implanted 005 glioma stem cells into *Rag1^-/-^*mice which lack T and B cells, and then individually transferred WT or *Ifng^-/-^* CD4^+^ or CD8^+^ T cells **(Figures S5E)**. Importantly, adoptive transfer of WT CD4^+^ T cells, but not CD8^+^ T cells or *Ifng^-/-^*CD4^+^ T cells into *Rag1^−/−^* mice led to MHC-II upregulation on microglia **(Figures S5F).** These intimate interactions between CD4^+^ T cells and microglia were directly confirmed by confocal microscopy that showed that most of the CD4^+^ T cells found in 005 gliomas after αCTLA-4 treatment are concentrated around ionized calcium-binding adaptor molecule 1 (Iba1^+^) and MHC-II^+^ microglia **(Figure 5G)**. In contrast, the IgG control gliomas displayed very little MHC- II-expressing microglia and the CD4^+^ T cells were more loosely scattered throughout the gliomas **(Figure 5G).** Lastly, in addition to enhancing MHC-II and antigen presentation on microglia, CD4^+^ T cells were also essential during αCTLA-4 treatment for maintaining microglial abundance **(Figure S5G)** and the recruitment of Ly6c^int^ monocytes and B220^+^ B cells while limiting the infiltration of Ly6c^hi^ monocytes and neutrophils into the brain **(Figures S5H-K).** Since depletion of B cells or monocytes (using *mu*MT^-^ or *Ccr2^-/-^* mice respectively) did not influence the therapeutic efficacy of αCTLA-4 **(Figures S3F-I)**, the role of these cells in the TME have yet to be determined. Altogether, these results indicated that IFNγ-producing CD4^+^ TILs remodel the TME and selectively develop a mutualistic partnership with microglia in gliomas wherein they support the generation of MHC-II^+^ antigen-presenting microglia that in return, sustain the anti-tumor Th1 functions.

### Microglial activation induced by CD4^+^ T cells promotes tumor phagocytosis via AXL/MER signaling

Given the pivotal role of CD4^+^ TILs and microglia in conferring protection against glioma, we next asked if the anti-tumor effects of CD4^+^ T cells depended on their direct recognition of the tumor cells or killing by IFNγ. We knocked out *Ciita* to disrupt MHC-II expression, even though MHC-II was virtually undetectable on 005 cells *in vivo* or *in vitro* (including after IFNγ stimulation (**Figures S6A**), or *Ifngr1* to disrupt IFNγ-signaling in 005 cells **(Figures S6A-C),** and examined if the tumors were sensitive or resistant to ICB. CIITA-deficient tumor cells were as sensitive to αCTLA-4 as scramble control tumor cells (**Figure 6A**), demonstrating that the protective effect of αCTLA-4 therapy did not require direct CD4^+^ T cell recognition of glioma cells via MHC-II. In contrast, IFNγR-deficient tumors were resistant to αCTLA-4 therapy, suggesting that the protective effects of αCTLA-4 were dependent on IFNγR signaling within the tumor cells **(Figure 6B)**. All together, these data suggest that while the CD4^+^ T cells were needed for ICB tumor regression, they did not kill the tumor cells through direct MHC-II-peptide recognition, but rather via IFNγ-mediated events.

**Figure 6.**
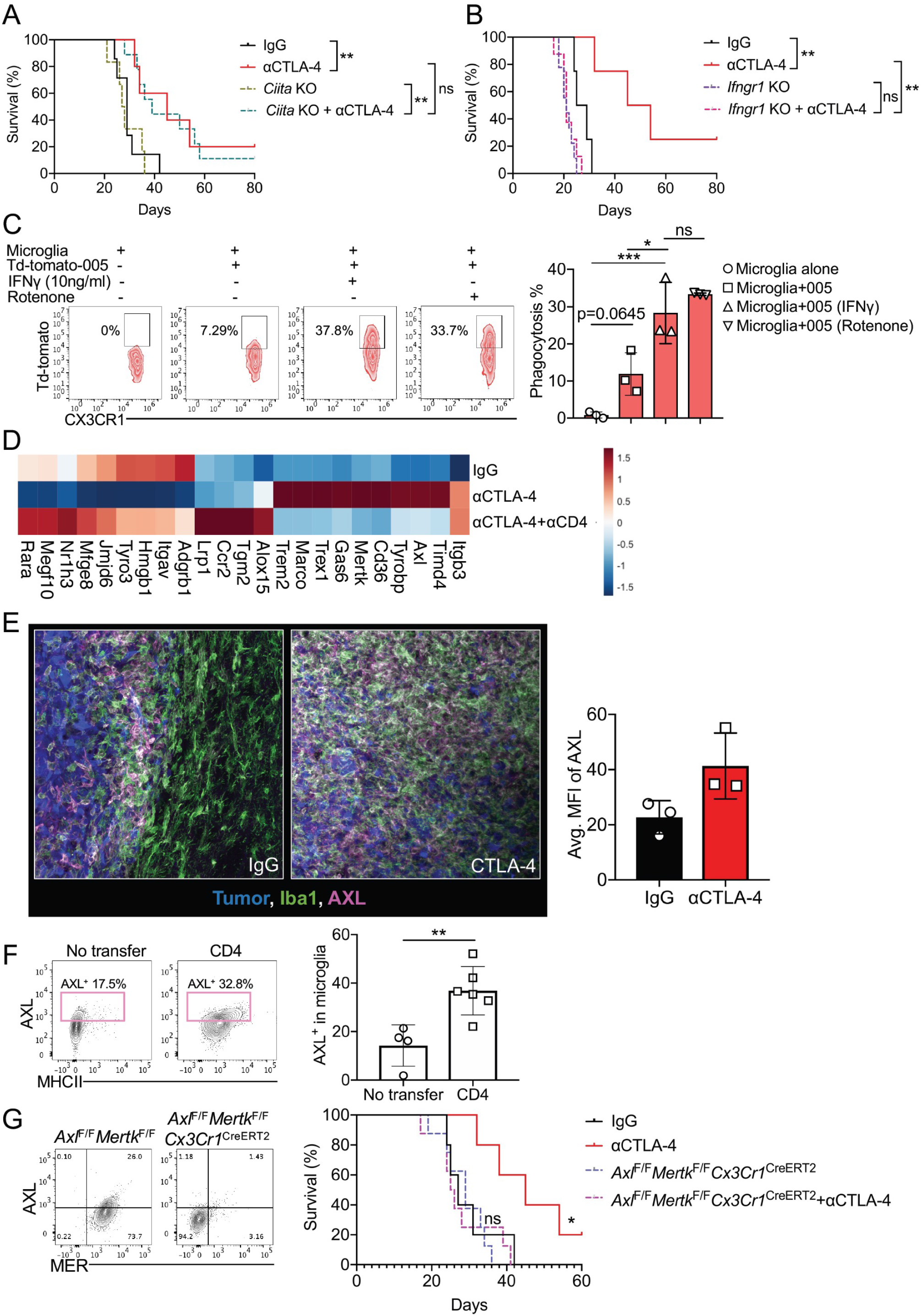
Tumoricidal activities of microglia are dependent on TAM receptor (AXL/MER). (A) Mice were implanted with 3 × 10^5^ 005 or *Ciita* KO 005 on day 0, treated with 10 mg/kg IgG or αCTLA-4 injected i.p. on days 7, 14, 21 and 28, and monitored for survival. Median survival of IgG (29 days; n = 7) was compared with αCTLA-4 (45 days; n = 5, p = 0.009). Median survival of *Ciita* KO (27.5 days, n=5) was compared with *Ciita* KO treated with αCTLA-4 (39 days; n = 9, p = 0.0085). Similarly, comparison of αCTLA-4 treatment of WT 005 with that of *Ciita* KO 005 (p = 0.876) were done by log-rank analysis. (B) Mice were implanted with 3 × 10^5^ 005 or *Ifngr1* KO 005 on day 0, treated with 10 mg/kg IgG or αCTLA-4 injected i.p. on days 7, 14, 21 and 28, and monitored for survival. Median survival of IgG (27 days; n = 4) was compared with αCTLA-4 (49.5 days; n = 4, p = 0.00673). Median survival of *Ifngr1* KO (21 days, n=9) was compared with *Ifngr1* KO treated with αCTLA-4 (21 days; n = 9, p = 0.4679). Similarly, comparison of αCTLA-4 treatment of WT 005 with that of *Ifngr1*KO 005 (p = 0.00213) were done by log-rank analysis. (C) tdTomato 005 GSCs cells were treated with IFNγ (10 ng/ml) or Rotenone (100 nM) for overnight. Microglia isolated from *Cx3cr1*^GFP^ mice were co-cultured with tdTomato 005 GSCs from different conditions *in-vitro* for 4 hours. tdTomato signal from microglia was measured by flow cytometry. (D) Heatmap showed the level of apoptotic cell clearance associated genes in microglia from bulk RNA-seq in different treatment among IgG vs. αCTLA-4 vs. αCTLA-4+ αCD4. (E) C57BL/6 mice were implanted with 3 × 10^5^ 005 GSCs on day 0, treated with 10 mg/kg αCTLA-4 or IgG injected i.p. on days 7, 14, 21, 28, and euthanized on day 35. The brains were harvested and fixed. The localization of Iba1^+^, AXL^+^ cells in the brain was analyzed by immunofluorescence confocal imaging. The average MFI of AXL was quantified. (F) Rag1^-/-^ recipient mice were transferred with or without CD4^+^ T cells (1x10^6^) cells on day 0. All Rag1^-/-^ mice were implanted with 3 × 10^5^ 005 GSCs on day 1. AXL expression on microglia within tumor tissue were analyzed and quantified on day 28. (G) *Axl^F/F^Mertk^F/F^Cx3cr1^CreERT^*mice were treated with 150 mg/kg tamoxifen for two doses 48 hours apart 1 week before, and then implanted with 3 × 10^5^ 005 GSCs on day 0, treated with 10 mg/kg IgG or αCTLA-4 on days 7, 14, 21 and 28, and monitored for survival. The efficacy of depletion of AXL and MER on microglia was checked by flow cytometry (left). Median survival of IgG (27 days; n = 5) was compared with αCTLA-4 (45 days; n = 5, p = 0.019). Median survival of *Axl^F/F^Mertk^F/F^Cx3cr1^CreERT^*(29 days, n=8) was compared with *Axl^F/F^Mertk^F/F^Cx3cr1^CreERT^*treated with αCTLA-4 (25.5 days; n = 9, p = 0.8502). Similarly, wild-type B6 mice treated with αCTLA-4 was compared with *Axl^F/F^Mertk^F/F^Cx3cr1^CreERT^* treated with αCTLA-4 (p = 0.016) by log-rank analysis (right). All results were pooled from or representative of 2-4 experiments with n = 3 mice / group (D), n = 3 mice / group (E), n = 4-6 mice / group (F). Data are expressed as mean ± SEM. Statistical analysis was performed using Student’s *t* test (two tailed) comparing IgG group to various treatment groups. * p < 0.05, ** p < 0.01, *** p < 0.001, **** p < 0.0001.

Although IFNγR expression on the GSCs was critical for the therapeutic efficacy of αCTLA-4, IFNγ did not appear to directly kill the tumor cells *in vitro* because culturing the 005 cells with IFNγ for 72 hours did not stall proliferation or growth, but it did induce the early apoptotic marker Annexin V (**Figure S6D**). Furthermore, the suppression of glioma growth was independent of CD8^+^ T cell detection (**Figure 1F**) because knocking out *B2m* on 005 cells also had no impact on αCTLA-4 therapy **(Figure S6E)**. Similarly, depletion of NKs also did not affect overall survival **(Figure S6F)**, suggesting that NK cells were not involved in anti-glioma immunity. Therefore, we reasoned that perhaps IFNγ−signaling makes 005 cells vulnerable to killing by another cell type such as microglia or macrophages. To test if IFNγ sensitizes tumor cells to be phagocytosed by microglia, 005-tdTomato cells were exposed to IFNγ and then the cells were cultured with microglia. Phagocytosis was read out as tdTomato^+^ microglia by flow cytometry. As a positive control, we cultured the microglia with tumor cells treated with rotenone (a mitochondrial poison) to induce tumor cell apoptosis. These experiments demonstrated that microglial phagocytosis of tumor cells was considerably elevated by treating the tumor cells with either IFNγ or rotenone **(Figure 6C)**, showing that IFNγ may sensitize tumor cells to be phagocytosed by microglia *in vitro*.

Additionally, comparison of DEGs and Gene Set Enrichment Analysis (GSEA) from bulk RNA-sequencing of microglia isolated from gliomas treated with IgG *vs.* αCTLA-4 *vs.* αCTLA-4 + αCD4 showed induction of genes involved in phagocytosis pathway after αCTLA-4 therapy that were dependent on CD4^+^ T cells **(Figure 6D and S6G)**. This included several genes involved in phagocytosis such as *Axl,* c-mer proto-oncogene tyrosine kinase (*Mertk*), growth arrest-specific gene 6 *(Gas6), Marco, Tyrobp, Cd36*. In particular, the receptor tyrosine kinase receptors *Axl*, *Mertk* of the Tyro3-AXL-MER (TAM) receptor family (Huang et al., 2021; Seitz et al., 2007) and their ligand *Gas6* were upregulated on microglia by αCTLA-4 therapy in a CD4^+^ T cell-dependent manner (**Figure 6D**). These receptors recognize and phagocytose apoptotic cells that expose phosphatidylserine (PtdSer), which was increased on 005 cells after IFNγ treatment *in vitro* (**Figure S6D**). Upon closer inspection, AXL was selectively induced in microglia surrounding the edges and within the tumor masses based on confocal microscopy, and this was further upregulated by αCTLA-4 therapy **(Figure 6E)**. Further, transfer 005 cells into *Rag1^-/-^* mice followed by the transfer of CD4^+^ T cells showed that AXL was preferentially upregulated on MHC-II^+^ microglia in a CD4^+^ T cell-dependent manner **(Figure 6F)**. Given that TAM receptors play an important role in phagocytosis of dying cells, we next asked if *Axl/Mertk*-deletion in microglia exert any impact on tumor regression during ICB. To do so, we utilized *Axl^F/F^Mertk^F/F^Cx3cr1^CreERT2^*mice to knock out *Axl* and *Mertk* in microglia (Huang et al., 2021). *Axl/Mertk*-deletion in microglia completely abrogated the therapeutic benefit of αCTLA-4 in glioblastoma **(Figure 6G)**, indicating that phagocytic function of microglia is critical for ICB-mediated tumor regression. In conclusion, these data show that GSCs can become sensitive to immune attack during ICB by creating a critical partnership between CD4^+^ T cells and microglia **(Figure S6H)**. Th1 CD4^+^ T cells are necessary to induce microglial antigen-presentation capacity via IFNγ-signaling that in turn, sustains (1) local CD4^+^ T cell effector functions, (2) potentiates the phagocytic functions of microglia via upregulation of phagocytic receptors like AXL/MER and (3) sensitizes tumor cells to immunosurveillance and removal by myeloid cells via increase in PtdSer. This microglia-CD4 T cell crosstalk highlights a novel therapeutic axis that enables robust protective anti-tumor immunity in preclinical models of glioblastoma and provides new insights for developing therapeutic strategies against human glioblastoma.

## Discussion

Despite several immunotherapy clinical trials with ICB, little-to-no clinical benefit has been observed in glioblastoma patients (Omuro et al., 2018). It remains to be determined if a protective immune response can be raised against glioblastoma, and if so, what is required to do so. Thus, using more physiological pre-clinical models of glioblastoma, we sought to develop a better understanding of the cellular and biological mechanisms that mediate immune responses to glioblastoma upon ICB treatment. Here we show that αCTLA-4 treatment significantly improved survival and suppressed the growth of 005 tumors that contain OPC- and MES-like GSCs. Surprisingly, the efficacy of αCTLA-4 was lost in mice that were depleted of CD4^+^ but not CD8^+^ T cells, suggesting that CD4^+^ T cells are the critical immune players that can infiltrate into the parenchyma of the brain, reprogram the TME and suppress GSC growth. Similarly, we found that microglia, the local sentinels of the CNS, also play a significant role in tumor control, as indicated by faster glioblastoma growth and reduced number of Th1 tumor-infiltrating CD4^+^ T cells when mice were depleted of microglia. Importantly, activation and expression of MHC-II on microglia was critically dependent on CD4^+^ but not CD8^+^ T cells, and microglial MHC-II expression was needed to sustain anti-tumor immunity by CD4^+^ T cells and suppression of tumor growth. Such a co-dependency between CD4^+^ T cells and microglia to target brain cancer has not been described before and therefore, this work highlights a fundamental axis that may lead to curative anti-tumor immune responses via ICB in human glioblastoma.

It is not clear why treatment with ICBs have failed in glioblastoma, but to date, most clinical trials have incorporated αPD-1 as opposed to αCTLA-4 ICB. A recent report by Arrieta *et al*. showed that a small subset of recurrent glioblastoma patients with high pERK1/2 responded to αPD-1 blockade and relevant to our study, these high pERK tumor cells are mainly colocalized with MHC-II^+^ microglia and other genes associated with antigen presentation (Arrieta et al., 2021). While this finding is suggestive of a role for microglia in tumor-antigen presentation, our report is the first to show that indeed, MHC-II^+^ microglia can present tumor antigens to intratumoral CD4^+^ T cells and sustain anti-tumor immunity to glioblastoma after ICB. αCTLA-4 induces the expansion of an ICOS^+^ Th1-like CD4^+^ T cells whereas αPD-1 blockade induces the expansion of effector CD8^+^ T cells (Ji et al., 2012; Ng Tang et al., 2013; Relecom et al., 2021; Wei et al., 2017). In support of our findings, the expression level of CTLA-4, but not PD-1, is associated with poor prognosis in glioblastoma patients (Liu et al., 2020). Our findings match clinical reports showing that decreased frequencies of anti-tumor CD4^+^ T cells in high-grade glioma patients are associated with adverse overall survival outcomes (Fecci et al., 2006; Grossman et al., 2011). In orthotopic glioma models, CD4^+^ CAR-T cells are effective in facilitating long-term anti-tumor response due to their outperformance over CD8^+^ CAR-T (Wang et al., 2018). Furthermore, a vaccine targeting mutant isocitrate dehydrogenase 1 (IDH1) is effective against IDH1^R132H^ glioma, which correlated with vaccine-induced IDH1^R132H^-specific CD40LG^+^ CXCL13^+^ CD4^+^ T cells but not CD8^+^ T cells (Platten et al., 2021; Wei et al., 2017). Our sc-RNAseq data also showed that both *Cd40lg* and *Cxcl13*, in addition to *Cxcr3* and *Cxcr6*, were upregulated in CD4^+^ population after αCTLA-4 treatment, in agreement with prior reports showing CD4^+^ T cells with these properties could traffic from the perivascular space to the parenchyma and provide anti-glioma protection (Platten et al., 2021; Wilson et al., 2010). Collectively, these findings demonstrate how CD4^+^ T cells can suppress glioblastoma in partnership with microglia, and therefore, we suggest that future clinical trials in glioblastoma should place more emphasis on αCTLA-4 ICBs alone or with other therapies that stimulate CD4^+^ T cells and microglia.

A general mechanism of resistance to ICB and CAR-T therapies is the loss of IFNγR signaling, including glioblastoma (Larson et al., 2022). Here, we found IFNγR signaling pathway in glioma was important for response to ICB. Loss of IFNγR signaling-related genes (IFNGR1, IFNGR2, JAK2) may serve as a biomarker to stratify glioblastoma patients for probability of response to ICB therapy. Interestingly, immune chemoattractant chemokine CXCL10, which binds to CXCR3 (Tokunaga et al., 2018), and fractalkine CX3CL1, which binds to CX3CR1, were upregulated in WT compared to *Ifngr1* KO gliomas from our bulk RNA-seq data (data not shown), which may explain enhanced immune cell recruitment and surveillance in WT vs. *Ifngr1* KO glioma. We also found IFNγ-signaling increased the apoptotic marker PtdSer on GSCs and increased their vulnerability to microglial phagocytosis, which aides in the clearance of apoptotic cells. TAM receptor recognition and engulfment of PtdSer-positive apoptotic cells are mediated by Gas6 and Protein S (Pros1) binding (Lemke, 2017; Lemke and Burstyn-Cohen, 2010; Scott et al., 2001), and further studies are required to elucidate whether microglial tumoricidal functions can be potentiated by increasing IFNγ signaling and AXL/MER activity in glioblastomas. As GAS6 and PROS1 are both highly expressed in MES-like glioblastoma (Hara et al., 2021), perhaps the IFNγ-PtdSer-AXL/MER axis could be potentiated by αCTLA-4 ICB to eradicate MES-like gliomas.

On this note, it may be more important to reprogram microglia than to deplete them for treating glioblastoma. Indeed, a phase II clinical trial of PLX3397 did not show significant efficacy in glioblastoma patients (Butowski et al., 2016), and similarly our data as well as that of (Rao et al., 2022) showed depletion of microglia actually promoted tumor growth. Why these results conflict with prior RCAS-hPDGF-B murine model (Pyonteck et al., 2013; Quail et al., 2016) is not entirely clear, but perhaps it has to do with GSC heterogeneity (Pyonteck et al., 2013; Yan et al., 2017). For example, the 005 model consists of OPC- and MES-like cells (Hara et al., 2021) and tumors progressed with CSF1R inhibitor (CSF1Ri) whereas the opposite was seen in the proneural-like PDGF-B-driven tumors (Herting et al., 2017; Quail et al., 2016; Rao et al., 2022). Further research is required to evaluate the response of CSF1Ri in various different glioma subtypes, which will better guide the application of CSF1Ri in human glioblastoma.

Lastly, our data demonstrate that CD4^+^ T cells, but not CD8^+^ T cells were uniquely situated to communicate with microglia via IFNγ-signaling for reasons that are still unclear. While both CD4^+^ and CD8^+^ T cells secrete IFNγ during glioblastoma pathogenesis, only IFNγ derived from CD4^+^ T cells led to MHC-II upregulation on microglia. One possible explanation for our findings is that there might exist quantitative differences in the amounts of IFNγ secreted from CD4^+^ versus CD8^+^ T cells that is not detected by intracellular cytokine staining. Another possibility is that CD4^+^ T cells may exist in closer spatial proximity and adhere more tightly to microglia than CD8^+^ T cells. A third possibility is that CD4^+^ T cells might have unique, unidentified features that are required to either activate microglia directly or recruit other cell types to stimulate microglia. Further work is needed to answer these questions. CD4^+^ T cells induced not only MHC-II expression on microglia, but also genes associated with antigen-presenting machinery (*Cd74, Cd86, Tap1*), and tumor sensing genes (*Gpr34, Spac*), and phagocytosis (*Axl, Mertk, Gas6, Marco, Tyrobp, Cd36*) on microglia. Consistent with our finding, a CD4^+^ T cell-microglia interaction has also been reported in neonatal development whereby microglia require CD4^+^ T cells to complete the fetal-to-adult maturation in the CNS (Locatelli and Engelhardt, 2020; Pasciuto et al., 2020). In summary, our study extends from the CD4^+^ T cell/microglia partnership by uncovering new ways in which such cooperation can be co-opted to develop more effective immunotherapies against human glioblastoma.

## Acknowledgements

We thank the Kaech lab and Dr. Youtong Huang and Dr. Kaisa Happonen for helpful discussion. 005 GSCs were kindly provided by Dr. Inder M. Verma. PLX3397 chow were kindly provided by Dr. Lara Labarta Bajo and Dr. Nicola Allen. CD169-DTR were kindly provided by Dr. Masato Tanaka. This work was supported by the Flow Cytometry Core Facility of the Salk Institute with funding from NIH-NCI CCSG: P30 014195 and Shared Instrumentation Grant S10-OD023689 (Aria Fusion cell sorter) and by the NGS Core Facility of the Salk Institute with funding from NIH-NCI CCSG: P30 014195, the Chapman Foundation and the Helmsley Charitable Trust. This work was supported by the NIH (NCI CA195613 to I.M.V. and T.H.), and Salk innovation grant (to S.M.K.), Salk Accelerators Fellowship Award (D.C.), the NOMIS Center (D.C., and S.K.V.), as well as the following postdoc fellowships from the Cancer Research Institute (S.K.V., and S.M.), Damon Runyon Cancer Research Foundation (T.M.), National Cancer Center fellowship (S.K.V.).

## Author Contributions

Conceptualization, D.C., S.K.V., S.M.K.; Methodology, D.C., S.K.V., T.H.; Formal Analysis, D.C., S.K.V., B.M.; Investigation, D.C., S.K.V., T.H., K.T., B.M., Y.F., J.C., S.X., Z.X., V.T., E. C., S.M., Q.Y., Y.Z., T.H., G.L.; Resources, C.O.; Writing – Original Draft, D.C.; Writing – Review & Editing, D.C., S.K.V., T.H., T.H.M., V.D., H.K.C., T.H., S.M.K.; Funding Acquisition, S.M.K.; Supervision, S.M.K.

## STAR METHODS

### KEY RESOURCES TABLE

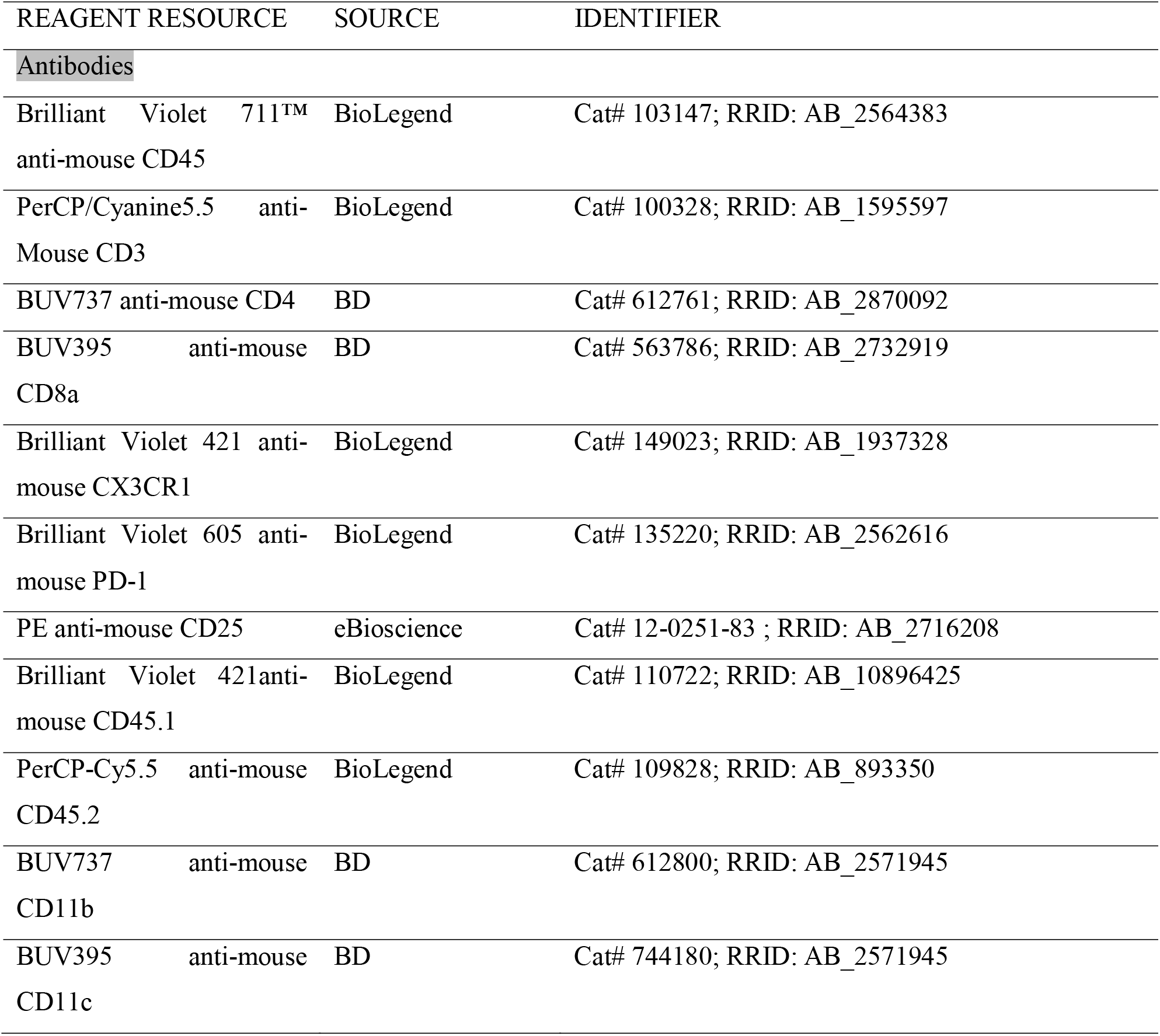
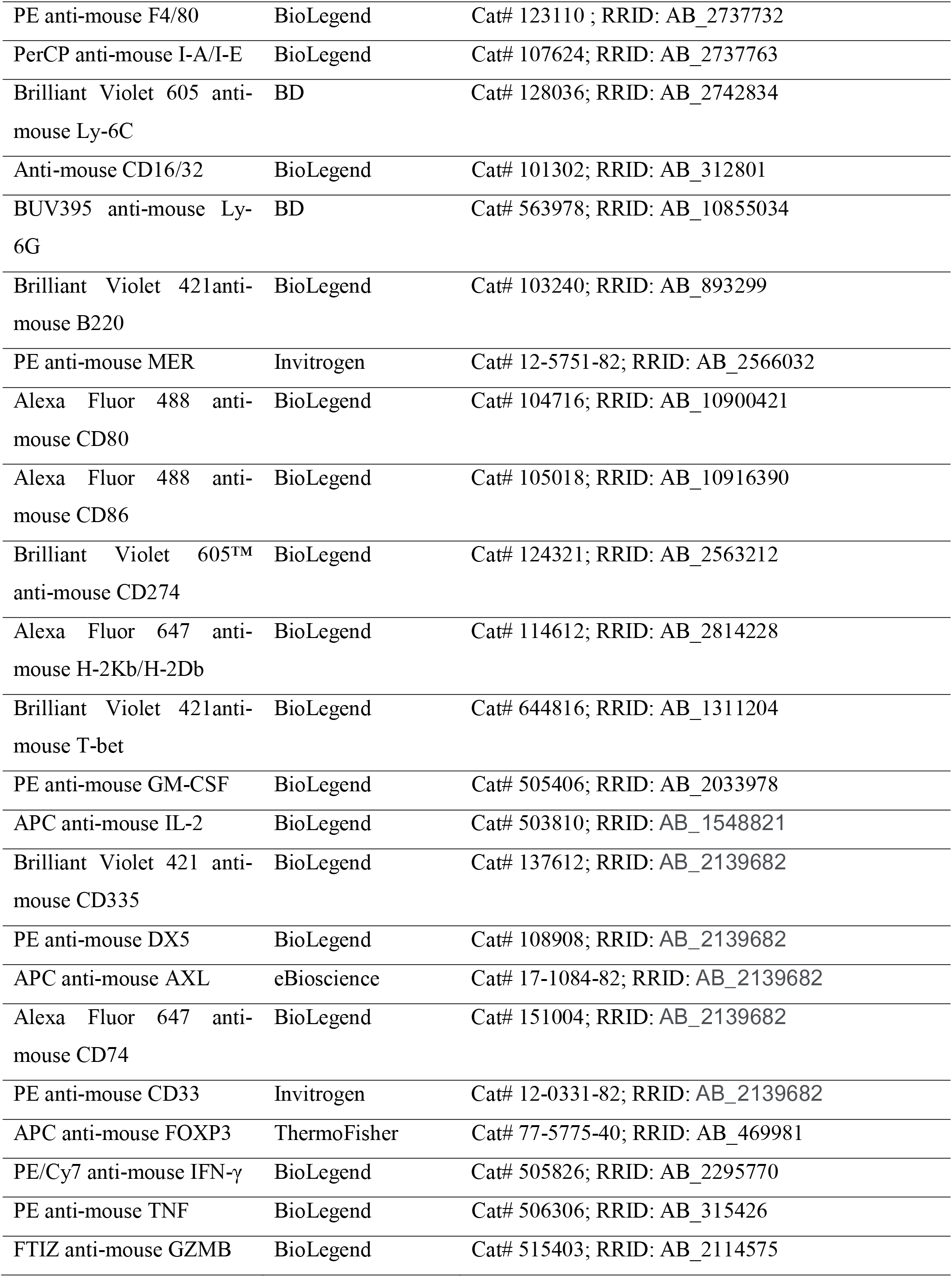
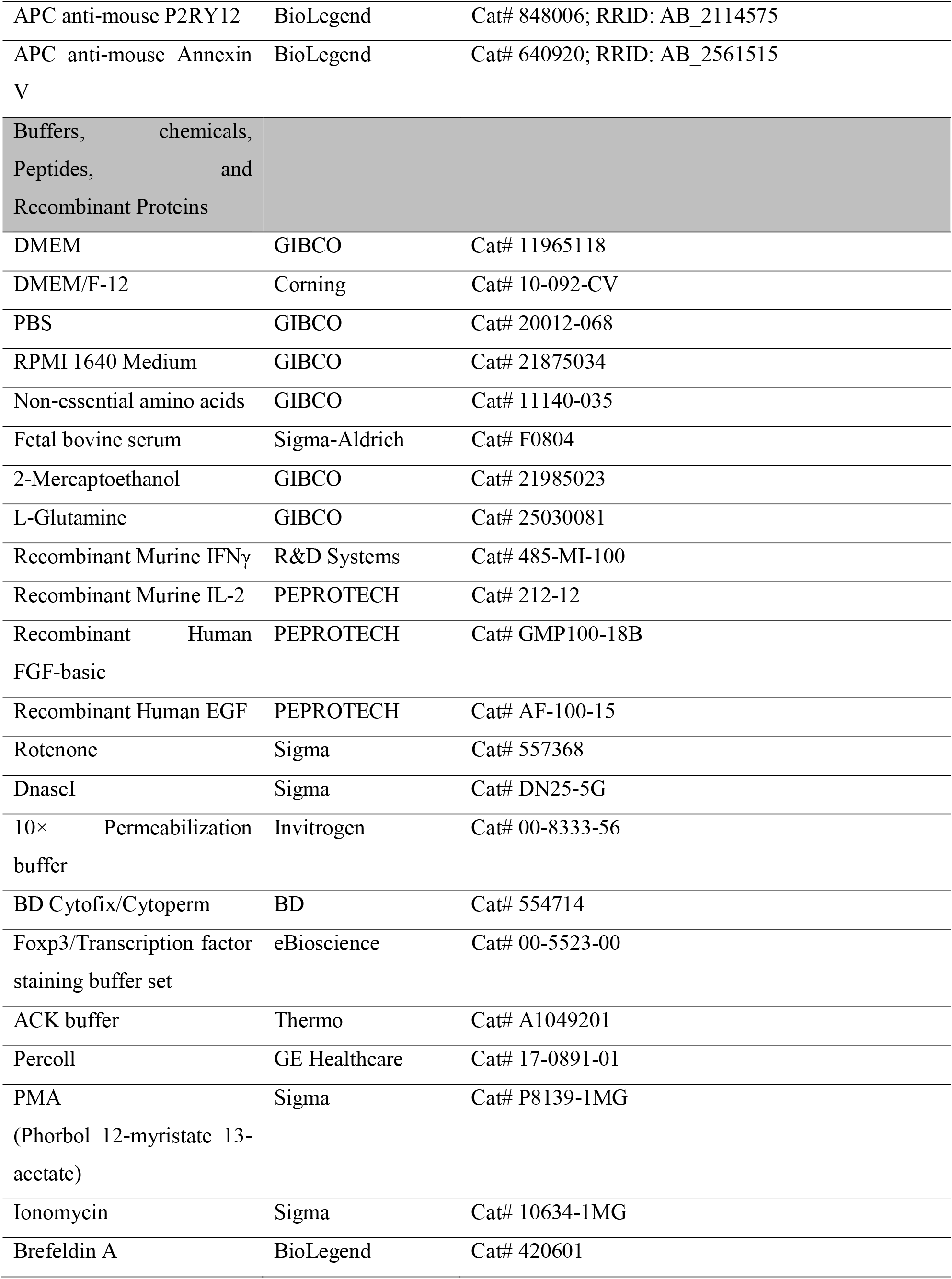
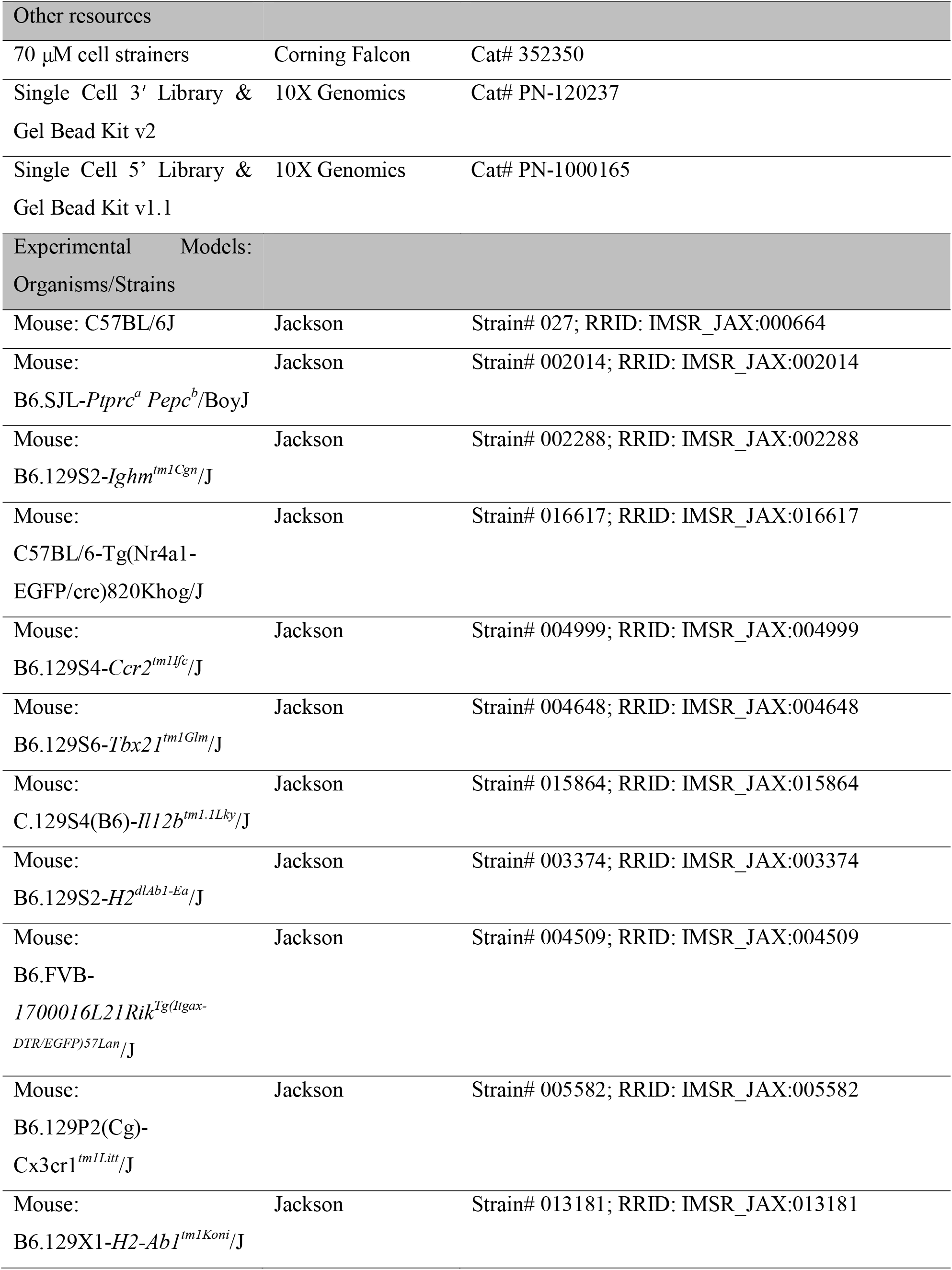
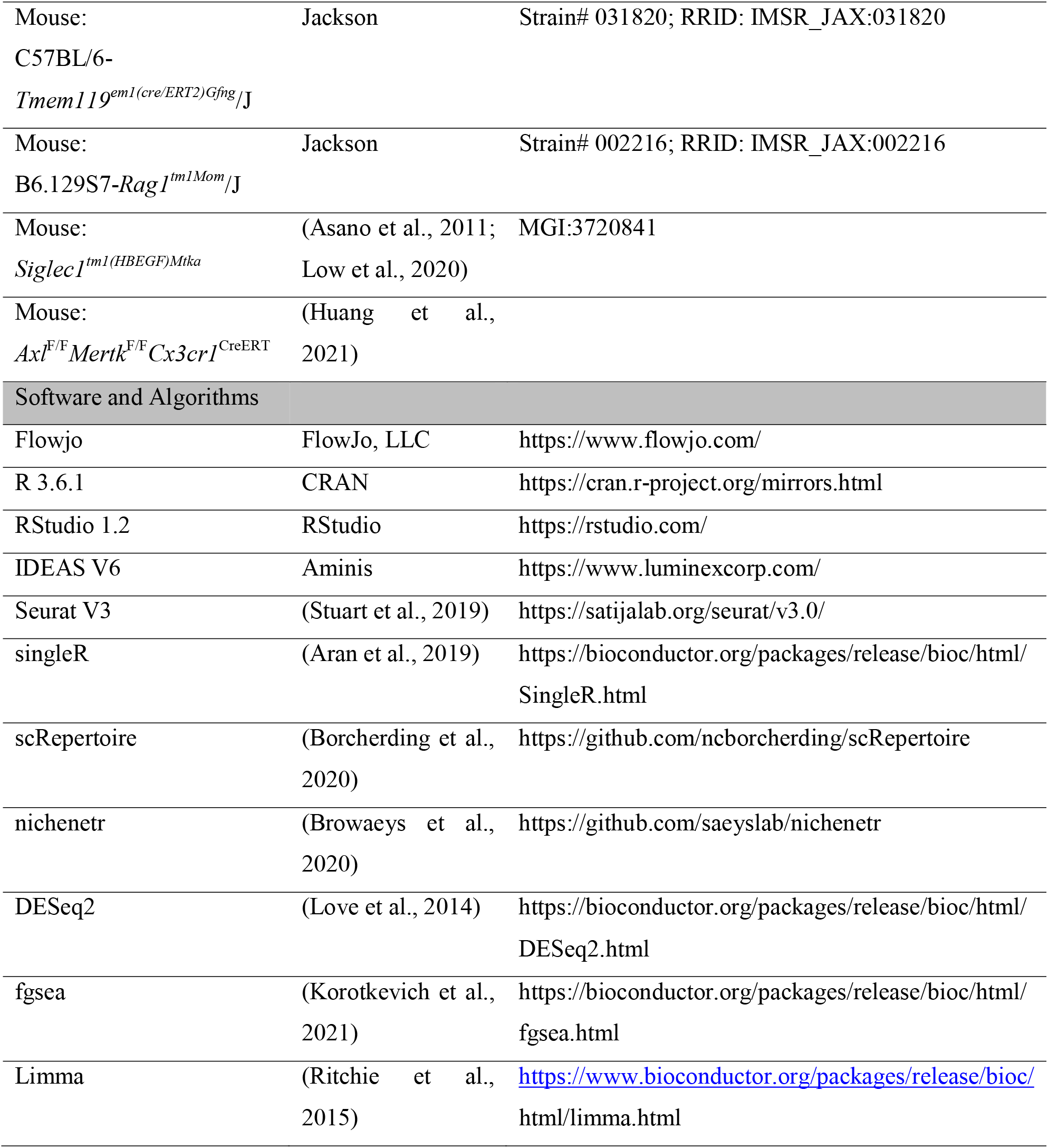

### RESOURCE AVAILABILITY

#### Lead Contact

Further information and requests for resources and reagents should be directed to and will be fulfilled by the lead contact, Susan M. Kaech.

#### Materials Availability

This study did not generate new unique reagents.

#### Data and Code Availability

All sequencing data from this paper will be available in the GEO database.

### EXPERIMENTAL MODEL AND SUBJECT DETAILS

#### Mice

B6 (C57BL/6J), Ly5.1 (B6.SJL-*Ptprc^a^ Pepc^b^*/BoyJ), *mu*MT^-^ (B6.129S2-*Ighm^tm1Cgn^*/J), Nur77^GFP^ (C57BL/6-Tg(Nr4a1-EGFP/cre)820Khog/J), *Ccr2^-/-^*(B6.129S4-*Ccr2^tm1Ifc^*/J), T-bet^-^ (B6.129S6- *Tbx21^tm1Glm^*/J), IL-12^YFP^ (C.129S4(B6)-*Il12b^tm1.1Lky^*/J), MHC-II^-^ (B6.129S2-*H2^dlAb1-Ea^*/J), CD11c- DTR (B6.FVB-*1700016L21Rik^Tg(Itgax-DTR/EGFP)57Lan^*/J), CX3CR1^GFP^ (B6.129P2(Cg)-

Cx3cr1*^tm1Litt^*/J), H2-Ab1-flox (B6.129X1-*H2-Ab1^tm1Koni^*/J), Tmem119-Cre ERT2 (C57BL/6- *Tmem119^em1(cre/ERT2)Gfng^*/J), and Rag1^-/-^ (B6.129S7-*Rag1^tm1Mom^*/J) mice were purchased from Jackson Laboratories. H2-Ab1^fl/fl^Tmem119^CreERT2/+^ mice were generated by crossing H2-Ab1- flox mice with Tmem119-Cre ERT2 mice. CD169-DTR (*Siglec1^tm1(HBEGF)Mtka^*) (Asano et al., 2011) were kindly provided by Dr. Masato Tanaka at the Tokyo University of Pharmacy and Life Sciences (Tokyo, Japan). *Axl*^F/F^*Mertk*^F/F^*Cx3ct1*^CreERT^ mice (Huang et al., 2021) were kindly provided by Dr. Greg Lemke at Salk Institute for Biological Studies (La Jolla, USA). All mice were maintained under specific-pathogen-free facilities at the Salk Institute and all experimental studies were approved and performed in accordance with guidelines and regulations implemented by the Salk Institute Animal Care and Use Committee.

#### Cell Lines

Mouse 005 glioma stem cells (GSCs) (Marumoto et al., 2009) or 005-Luciferase (005-Luc) GSCs were cultured as spheres in serum free stem cell medium composed of advanced DMEM/F12 medium (Corning), supplemented with 2 mM L-glutamine (Corning), 1% penicillin-streptomycin (Corning), 1% N2 supplement (Gibco), 40 mg/mL heparin (Sigma-Aldrich), recombinant human EGF (20 ng/mL; PeproTech), and recombinant human FGF-basic (20 ng/mL; PeproTech). Spheres were dissociated with TrypLE™ Express (ThermoFisher) for passaging. All the rest of the genetically-modified cell lines, including *Ciita* KO 005 GSCs, β*2m* KO 005 GSCs, *Ifngr1* KO 005 GSCs, and tdTomato overexpression (OE) 005 GSCs were all generated based on 005 GSCs cells. GL261 glioma cell line was a gift from Jeremy N. Rich (University of Pittsburgh) and was cultured in Dulbecco’s Modified Eagle Medium (DMEM) (ThermoFisher) with 10% fetal bovine serum and 1% penicillin-streptomycin. Cells were dissociated with 0.25% Trypsin-EDTA (ThermoFisher). Both 005 GSCs and GL261 were cultured at 37 °C in 5% CO_2_.

#### Intracranial Glioma Mouse Model

Single cell suspension of glioma cell lines was stereotactically injected into the hippocampus of 8- to 12-week-old C57BL/6J mice or other mice stains mentioned in the Mice Section above. Briefly, 005 GSCs (3×10^5^ cells) or GL261 (5×10^4^ cells) suspended in 3 μL PBS were loaded on a Hamilton microliter syringe with 26-gauge needle, and injected at a speed of 0.1uL/30s-1min using the following coordinate: 2.0 mm posterior, 1.5 mm lateral, and 2.3 mm dorsal to the bregma. Upon completing injection, the needle was left in place for 3 min, then withdrawn slowly in 2 min to help reduce cells reflux. 7 days after tumor implantation, mice were randomly divided into groups and injected with antibodies αPD-1 (clone RMP1-14; 10 mg/kg), or αCTLA- 4 (Syrian hamster clone 9H10; 10 mg/kg). Isotype control antibodies rat IgG2a (clone LTF-2) or Syrian hamster IgG were purchased from BioXcell and administered 4 times intraperitoneally (i.p.). For depletion/blockade studies, C57Bl/6 mice were implanted with 005 GSCs and 7 days after implantation mice were treated with depletion antibodies aCD4 (clone GK1.5), aCD8 (2.43), NK1.1 (clone PK136), IFNγ blockade (clone XMG1.2) or rat IgG at 10 mg/kg. Mice were followed for neurological symptoms and euthanized before becoming moribund.

### METHOD DETAILS

#### Tumor digestion and cell isolation

All mice were perfused with ice-cold PBS after euthanasia to reduce the influence of peripheral blood derived cells. All the brains were collected in ice-cold PBS, and then mechanically and enzymatically dissociated using a papain-based neural tissue dissociation kit P (Miltenyi Biotec) supplemented with 0.1% type I collagenase (Thermo Fisher Scientific) in a C-tube (Miltenyi Biotec), and then dissociated with a gentleMACS Qcto Dissociator using program 37C_NTDK_1 for 30 min, followed by filtration with 70 µm cell strainers. Filtered cells were incubated with ACK lysis buffer (Invitrogen) to lyse red blood cells. For glioma single cell suspension, filtered cells were resuspended with 0.9 M sucrose, spun down to remove supernatant, and the pellet containing tumor cells was saved for later use. For immune cell suspension, filtered cells were resuspended in 5 mL 37% Percoll, and slowly added beneath the 70% Percoll, and then centrifuged with brakes off. The immune cells were isolated from the interface of 37% and 70% Percoll and transferred into excessive RPMI 1640 medium containing 10% fetal bovine serum for later use.

#### Flow cytometry, cell sorting and antibodies

Both single cell suspensions were incubated with Fc receptor-blocking anti-CD16/32 (BioLegend) on ice for 10 min before staining. Cell suspensions were first stained with Red Dead Cell Stain Kit (ThermoFisher) for 10 min on ice. Surface proteins were then stained in FACS buffer (PBS containing 2% FBS and 0.1% sodium azide) for 30 min at 4°C. To detect cytokine production *ex-vivo*, cell suspensions were re-suspended in RPMI 1640 containing 10% FBS, and then stimulated by 50 ng/ml PMA and 3 μM Ionomycin in the presence of 2.5 μg/ml Brefeldin A (BioLegend #420601) for 4 h at 37°C. Cells were processed for surface marker staining as described above. For intracellular cytokine staining, cells were fixed in BD Cytofix/Cytoperm (BD #554714) for 30 min at 4 °C, then washed with 1× Permeabilization buffer (Invitrogen #00-8333-56). For transcription factor staining, cells were fixed in Foxp3 / Transcription Factor Fixation/Permeabilization buffer (Invitrogen #00-5521-00) for 30 min at 4 °C, then washed with 1× Permeabilization buffer. Cells were then stained with intracellular antibodies for 30 min at 4 °C. Samples were processed on LSR-II flow cytometer (BD Biosciences) and data were analyzed with FlowJo V10 (TreeStar). For sorting experiments, cells were sorted either on FACSAria™ III sorter or Fusion sorter (BD Biosciences). The following antibodies against mouse proteins were used: anti-CD45 (30-F11), anti-CD3ε (145-2C11), anti- CD4 (GK1.5), anti-CD8a (53-6.7), anti-PD-1 (29F.1A12), anti-CX3CR1 (SA011F11), anti- Ly5.1 (A20), anti-Ly5.2 (104), anti-CD25 (PC61.5), anti-CD49b (DX5), anti-NKp46 (29A1.4), anti-I-A/I-E (MHC-II) (M5/114.15.2), anti-CD80 (16-10A1), anti-CD86 (GL-1), anti-PD-L1 (10F.9G2), anti-CD11b (M1/170), anti-CD11c (N418), anti-CD169 (3D6.112), anti-F4/80 (BM8), anti-Ly6c (HK1.4), anti-Ly6G (1A8), anti-B220 (RA3-6B2), anti-CD19 (1D3/CD19), anti-MER (DS5MMER), anti-CD74 (In1/CD74), anti-AXL (175128), anti-P2RY12 (S16007D), anti-H-2Kb/H-2Db (MHC-I) (28-8-6), anti-FoxP3 (FJK-16S), anti-IFN-γ (XMG1.2), anti-TNF-α (MP6-XT22), anti-GZMB (GB11), anti-IL-2 (JES6-5H4), anti-GM-CSF (MP1-22E9), and anti-T-bet (4B10). These antibodies were purchased from Invitrogen, BioLegend, Cell Signaling, or eBiosciences.

#### Tissue sectioning and Immunohistochemistry

Mice were euthanized and transcardially perfused with PBS followed by 2% paraformaldehyde (PFA). Dissected brains were fixed in 2% PFA overnight at 4°C followed by ten-minute washes for three times in PBS. Brains were mounted for sectioning using 4% low melting point agarose (Sigma, A9414) dissolved in water. Serial coronal sections (75µm) were taken using a vibratome (Leica VT1000S). Brain sections were incubated in blocking buffer (1× PBS, 0.03% Triton X- 100, 0.015% Tween20, 2% BSA) for 2 hours at room temperature. Primary antibody incubations were done in blocking buffer for 2 days at 4°C on a shaker. Excess primary antibodies were rinsed out by thirty-minute washes for three times with wash buffer (1× PBS, 0.03% Triton X100, 0.015% Tween20). Secondary antibody incubations were done overnight at 4°C. Excess antibodies were removed by thirty-minute washes for three times with wash buffer. After labeling, all sections were mounted in several drops of Fluormount (Sigma, F4680) on standard 75mm x 25mm microscope slides (Corning, 2947) and sealed with a #1 coverslip and nail polish. The following antibodies against mouse proteins were used: Goat anti mouse AXL (AF854, R&D Systems), Rabbit anti mouse Iba1 (A19776, AB clonal), Anti-mouse CD4-PE (100407, Biolegend), Anti-mouse I-A/I-E (MHC Class II)-BV421 (107631, BioLegend), Donkey anti goat IgG AF594 (A-11058, Thermo Scientific), and Donkey anti rabbit IgG AF647 (A-31573, Thermo Scientific).

#### Microscopy and quantification

Brain sections were imaged with a Zeiss LSM 880 with a 20X air objective. Zeiss Zen Black software was used to set up confocal imaging parameters within the “Smart Setup” (“Best Signal” mode) feature based on the fluorophores used in immunofluorescence experiments. AXL expression was quantified from 4 to 5 sections per brain taken at least 150 µm apart. AXL fluorescence was measured from the tenth optical section within the z-stack of the tumor area from the brain section. All measurements of expression were done in Fiji (ImageJ) (Shihan et al., 2021). Mean fluorescence intensity (MFI) was calculated for the optical sections via Fiji (Image J) by calculating mean fluorescence on the AXL^+^ region of interest (ROI). Within the “Set Measurements” menu of the Fiji Results tab, we selected area, mean gray value, integrated density, and limit to threshold. The MFI of AXL expression was taken by setting an ROI from within the tumor border using the rectangle tool in Fiji from the 594nm detection channel used to image AXL expression. Background fluorescence was measured from an ROI outside the tumor border lacking AXL^+^ microglia. The MFI of the section was corrected by subtracting the MFI of the background ROI from the MFI of the AXL^+^ ROI within the tumor. The average MFI was determined by averaging corrected MFI measurements among the optical sections of a given brain. Statistical analysis of AXL expression was performed using GraphPad Prism (v9.3). Representative confocal images of tumors and associated immune cells were deconvoluted and prepared using Imaris (v9.8).

#### IVIS imaging

Glioma bearing mice were injected i.p. with 150 mg/kg D-Luciferin Firefly (Biosynth Carbosynth) in PBS. After 10 min the mice were anaesthetized with 2% isoflurane, and were imaged using the IVIS Spectrum In Vivo Imaging System (Xenogen).

#### Brain-protected bone marrow chimeric model

Ly5.1 or MHC-II^-^ (Ly5.2) recipient mice were irradiated at 1200 rads with lead shield brain- protection and reconstituted with 5 million bone marrow cells per mouse isolated from tibias and femurs of Ly5.1 donor mice. After ten weeks of reconstitution, immune chimeras were generated as previously described (Low et al., 2020).

#### CRISPR/Cas9 knock out in primary T cells and Adoptive T cell transfer

Naïve T cells were harvested from spleen and LNs. Naive CD4^+^ or CD8^+^ T cells were enriched using the EasySep Mouse CD4^+^/CD8^+^ T Cell Isolation Kit (STEMCELL Technologies) according to the manufacturer’s protocol. Cells were counted and resuspended at 5×10^6^ cells in PBS in a 1.5 mL Eppendorf tube per sgRNA target. For sgRNA/Cas9 ribonucleoprotein (RNP) complex formation, 0.6 µl of Cas9 protein (IDT) was combined with 1 µl of target sgRNA1, 1 µl of target sgRNA2 (0.3 nM, Synthego) and 2.4 µl of nuclease free water (CRISPRevolution kit) to a final volume of 5 µl in PCR tubes, and the mixture was incubated for 10 minutes at room temperature for complex formation. After complex formation, T cells were spun down, with supernatant removed from cell pellet. T cells were then resuspended in 20 μL of P3 buffer (16.4 μL P3 + 3.6 μL regent 1) (P3 Primary Cell 4D-Nucleofector™ X Kit S), with addition of 5 μl of RNP followed by gentle mixing. 25 μL of RNP/cell mix were then transferred to the bottom hole of a well of a Lonza nucleofector strip. Cells were electroporated by Lonza nucleofector using a pre-established DN100 program. Immediately after electroporation, 150 μL of 37 °C pre-warmed 10% RPMI were added and cells were let to rest for 10 minutes in the incubator. 1×10^6^ CD4^+^ or CD8^+^ cells per mice were adoptively transferred to Rag1^-/-^ mice 1 day before glioma implantation. The sequences of sgRNA are shown below: *Ifng* sgRNA1 (UCUAUGCCACUUGAGUUCUG), *Ifng* sgRNA1 (UCAAGACUUCAAAGAGUCUG); *Ciita* sgRNA1 (AGCUCGACUAAGGCUCCGGG), *Ciita* sgRNA2 (UCCAGUGUCCUAAUCUACCA); *B2m* sgRNA1 (AUUUGGAUUUCAAUGUGAGG), *B2m* sgRNA2 (ACUCACUCUGGAUAGCAUAC). *Ifngr1*sgRNA1 (UAUGUGGAGCAUAACCGGAG), *Ifngr1* sgRNA2 (GGUAUUCCCAGCAUACGACA).

#### Microglia isolation

Primary microglia were isolated by using Neural Tissue Dissociation Kit (P) (Miltenybiotec # 130-092-628) according to the manufacturer’s protocol. Briefly, brain tissues were collected from CX3CR1^GFP^ (B6.129P2(Cg)-Cx3cr1*^tm1Litt^*/J) mice, and they were washed in cold Dulbecco’s phopshate-buffered saline (D-PBS). Brains were then cut into small pieces and transferred into C tubes (Miltenybiotec # 130-093-237) containing 1900 μL buffer X, 50 μL enzyme P, 20 μL buffer Y and 10 μL enzyme A, and then dissociated with gentleMACS Program 37C_ABDK_01. After termination of the program, samples were resuspended and strained through 70 μm nylon strainer placed on a 50 mL tube. Filtered cells were centrifuged at 300×g for 5 minutes at 4 °C, and were resuspended in 5 mL 37% Percoll. These resuspended cells were slowly added beneath the 70% Percoll, and then centrifuged with brakes off. The immune cells were isolated from the interface of 37% and 70% Percoll and transferred into DMEM medium containing 10% fetal bovine serum for later use.

#### TCGA Survival analysis

RTCGA was used to obtain glioblastoma patients survival statistics and matched RNA expression values. The ratio of expression values for P2RY12 divided by CD163 was calculated for each patient. Patient samples were categorized into either high (greater than 50th percentile) or low (lower than 50th percentile) groups according to the P2RY12/CD163 ratio, and survival curves were plotted using the ggplot function in the Seurat package in R.

#### Single-cell RNA sequencing

Microglia (CD45^int^CX3CR1^+^), myeloid cells (CD45^hi^CD11b^+^), T cells (CD45^hi^CD3^+^) and all remaining immune cells (CD45^hi^CD11b^-^CD3^-^) were sorted from normal brain and 005 glioma-bearing brains treated with IgG, anti-CTLA-4, or αCTLA4 combined with CD4 depletion. Cells from each treatment group were pooled at equal ratios and then subjected to scRNA-seq. Sorted cells were partitioned into an emulsion of nanoliter-sized droplets using a 10x Genomics Chromium Single Cell Controller and RNA sequencing libraries were constructed using the Chromium Single Cell 3′ Library & Gel Bead Kit v2 (10X Genomics, Cat# PN-120237). Briefly, droplets containing individual cells, reverse transcription reagents and a gel bead were loaded with poly(dT) primers that include a 16 base cell barcode and a 10-base unique molecular index (UMI). Reverse transcription reactions were engaged to generate barcoded full-length cDNA followed by the disruption of emulsions using the recovery agent and cDNA clean up with DynaBeads MyOne Silane Beads (Thermo Fisher Scientific, Cat# 37002D). Bulk cDNA was amplified, and indexed sequencing libraries were constructed using the reagents from the Chromium Single Cell 3′ v2 Reagent Kit. Libraries were sequenced on NextSeq 500 Sequencing System (Illumina Cambridge).

#### Single-cell V(D)J sequencing

CD3^+^ cells were sorted and immediately loaded onto a 10x Chromium Chip G using reagents from the Chromium Single-Cell 5’ Library and Gel Bead Kit v1.1 (10x Genomics) according to the manufacturer’s protocol. 5’ gene expression libraries were generated using 10X Genomics Chromium Single Cell 5’ Library Construction Kits according to manufacturer protocols. TCR enrichment libraries were prepared following mouse T cell Single Cell V(D)J Enrichment Kits (10x Genomics). The final libraries were profiled using the Bioanalyzer High Sensitivity DNA Kit (Agilent Technologies) and quantified using the Kapa Library Quantification Kit (Kapa Biosystems). Each single-cell RNA-seq library was sequenced in one lane of NovaSeq6000 (Illumina) to obtain a minimum of 20,000 paired-end reads (26L×L91Lbp) per cell. Single-cell TCR V (D)J libraries were multiplexed and sequenced in one lane of NovaSeq6000 (Illumina) to obtain minimum of 5000 paired-end reads (26L×L91Lbp) per cell. The sequencing specifications for both single-cell RNA-seq and TCR V(D)J libraries were determined according to the manufacturer’s specification (10x Genomics).

#### Bulk RNA sequencing

Microglia (CD45^int^CX3CR1^+^) were sorted from the glioma bearing mice in different treatment groups (IgG vs. αCTLA-4 vs. αCTLA-4+αCD4). Total RNA was isolated using Trizol–Qiagen RNAeasy Micro kit and sent to Salk next generation sequencing core for ultra-low input RNA-Seq cDNA library preparation and sequencing run on Illumina Hiseq2500.

#### Bulk RNA-seq analysis

RNA sequencing was performed using the Illumina HiSeq 2500 platform on 51-bp single-end libraries prepared using the IlluminaTruSeq RNA sample preparation kit. Reads were mapped and aligned using STAR to the Mus musculus genome mm10. Reads were quantified and normalized using HOMER to generate raw counts and fragments per kilobase of exon per million mapped fragments (FPKM). Resulting FPKM values were processed by DESeq2 package (Love et al., 2014) to perform statistical analysis (Log2Fold change and p value). To identify enriched pathways, Gene Set Enrichment Analysis (GSEA) was performed using fgsea package (Korotkevich et al., 2021) on differentially expressed genes with L2FC>1.5 and p value<0.05.

#### Processing of scRNA/scTCR seq data

Raw sequencing data was processed and aligned mm10 mouse reference genome with CellRanger (10x Genomics) v 3.0.1. Resulting filtrated matrices (count matrices) of molecular counts were used as input for further processing with Seurat package V3.0.3 (Stuart et al., 2019) running under R studio. First, quality control was performed to create Seurat object with min features >200 and removal of cells having <200 or >5000 expressed genes or >7% mitochondrial counts. Variable features using FindVariableFeatures (using RNA and vst as an assay and selection method as parameters) and normalization using scTransform functions were performed. Further sc-transformed objects were integrated using PrepSCTIntegration, FindIntegrationAnchors and IntegrateData with normalization method set to “SCT”. This integrated data is scaled using ScaleData function using all genes and “RNA” as assay method. Principal Component Analysis was performed on top variable features using RunPCA and first 30 Principal components were chosen for computing shared nearest neighbors (SNN) using FindNeighbors function. FindClusters function with resolutions obtained from 0 to 1 in gradients of 0.1 was applied to our analysis. Finally, non-linear dimensional reduction such as UMAP/tSNE analysis was performed to visualize the data using RunUMAP and RunTSNE functions, respectively. To identify cell types, FindAllMarkers function was performed to determine differentially expressed features (cluster specific makers) and cluster names were manually curated. This was later confirmed using singleR package (Aran et al., 2019) and singlecellexperiment function. Top 5 genes from each cluster were used to further generate heatmap using DoHeatmap function. For each cluster, all differentially expressed genes (DEGs) were obtained by comparing different conditions using FindMarkers function (with assay = “RNA” and logfc.threshold = 0), which performs Wilcoxon Rank Sum test. This table is further used to generate volcano plot using ggplot package in Seurat. For plotting specific genes of interest, either DoHeatmap or DittoHeatmap function of dittoseq package is used (assay = “RNA”). Specific gene of interest was plotted on the UMAP using FeaturePlot package (slot = “scaled.data”) within Seurat.

#### NicheNet analysis

Cell interactions NicheNet Ligand-receptor interactions between populations from scRNA dataset were performed using nichenetr package (Browaeys et al., 2020). Required data that include NicheNet networks and ligand-target matrix were loaded. Ableit ligand target matrix was modified to include mouse antigen presenting genes. Receiver was set to microglial cluster, in order to determine the cell type contributing to the gene expression changes in microglia. Clusters representing CD8^+^ T cells, CD4^+^ helper T cells, Myeloid cells, Tregs and others are considered as sender cells. Nichenet analysis was performed as described in (https://github.com/saeyslab/nichenetr/blob/master/vignettes/seurat_wrapper_circos.md). The analysis was carried out especially on differentially expressed genes in microglia compared between anti-CTLA4 and IgG using FindMarkers function of Seurat package (with min.pct = 0.30 and assay = “RNA”). Finally, Circos plots using circos.par, chordDiagram, circos.track functions of circlize package were used to plot the ligand-receptor interactions.

#### scTCR seq analysis

scTCR and scRNA seq analysis was performed by combining Seurat and scRepertoire packages (Borcherding et al., 2020). For reading scTCR seq data, Read10X_h5 function of Seurat package was used and clonotype information was added to the single cell object using add_clonotype function. Following similar data processing as described above and upon removing cells having <200 or > 4000 expressed genes or >10% mitochondrial counts, objects were integrated, normalized (SCTransform) and scaled to determine principal components and SNN. TCR information (after combining filtered contig annotations of IgG and anti-CTLA4) was merged to integrated Seurat object using combineExpression function of scRepertoire package. Information of clonal expansion for each cell was determined and added to the Seurat object. Thus, “Hyperexpanded” cells are cells with clones between 100 – 500. Based on the number of clones, remaining cells were characterized as “Large” (20- 100), “Medium” (5 - 20), “Small” (1 - 5), or “Single” (0 -1). Cells with no TCR information are labelled as “NA”. Clonotypes within CD4 T cells/CD8 T cells/Treg were separately compared between IgG and anti-CTLA4 using alluvialClonotypes function. Specific TCR gene or amino acid sequence or non-TCR gene was plotted on the UMAP using highlightClonotypes function from scRepertorire package or FeaturePlot package of Seurat.

### QUANTIFICATION AND STATISTICAL ANALYSIS

Statistical analyses were performed using the two-tailed, unpaired, Student’s t-test unless otherwise specified. Each point represented a biological replicate and all data were presented as the mean ± SEM. The *P* values were represented as follows: ns, not significant, \*\*\*\**P* < 0.0001, \*\*\**P* < 0.001, \*\**P* < 0.01 and \**P* < 0.05.

## Supplemental information titles

**Figure S1.**
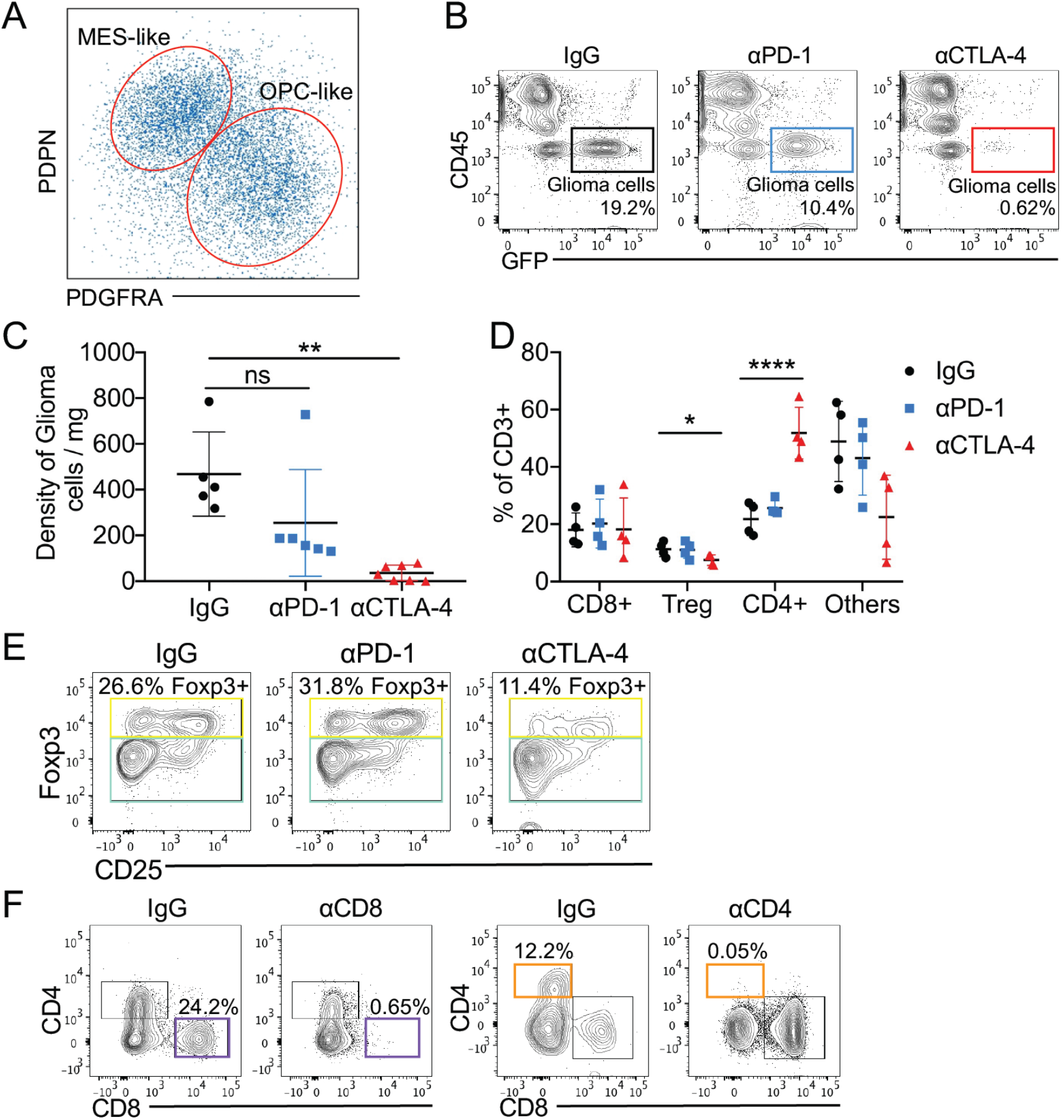
αCTLA-4 treatment increased the ratio of Non-treg/Treg, and suppressed tumor growth in glioblastoma bearing mice model. (A) Representative flow plots of MES-like marker PDPN against OPC-like marker PDGFRA on the glioblastoma cells in vivo from 005-bearing mice. (B-C) 005 GSCs (3 × 10^5^) were implanted on day 0, and treated with IgG, αCTLA-4, or αPD-1 on days 7, 14, 21 and 28, with euthanasia on day 35. Brain tissues were harvested, and glioma cells’ frequency (CD45^-^GFP^+^) (B) and density (C) in brain tumor tissues were measured and quantified by flow cytometry. (D) Frequency of CD8^+^ T cells (CD3^+^CD8^+^), non-Treg CD4^+^ T cells (CD3^+^CD4^+^Foxp3^-^), Treg cells (CD3^+^CD4^+^Foxp3^+^), and Others (CD3^+^CD4^-^CD8^-^) in brain tumor tissues from various treatment groups were measured and quantified by flow cytometry. (E) Representative flow plots of the frequency of Treg cells (% of FOXP3^+^ cells in CD4^+^ T cells) and non-Treg cells (% of FOXP3^-^ cells in CD4^+^ T cells) after different treatment from tumor tissue. (F) Representative flow plots of CD4^+^ or CD8^+^ T cells depletion in brain and spleen. All results were pooled from or representative of 2-4 experiments with n = 5-7 mice / group (B-C), n = 4 mice / group (D). Data are mean ± SEM. * p < 0.05, ** p < 0.01, *** p < 0.001, **** p < 0.0001 (two tailed unpaired Student’s *t* test).

**Figure S2.**
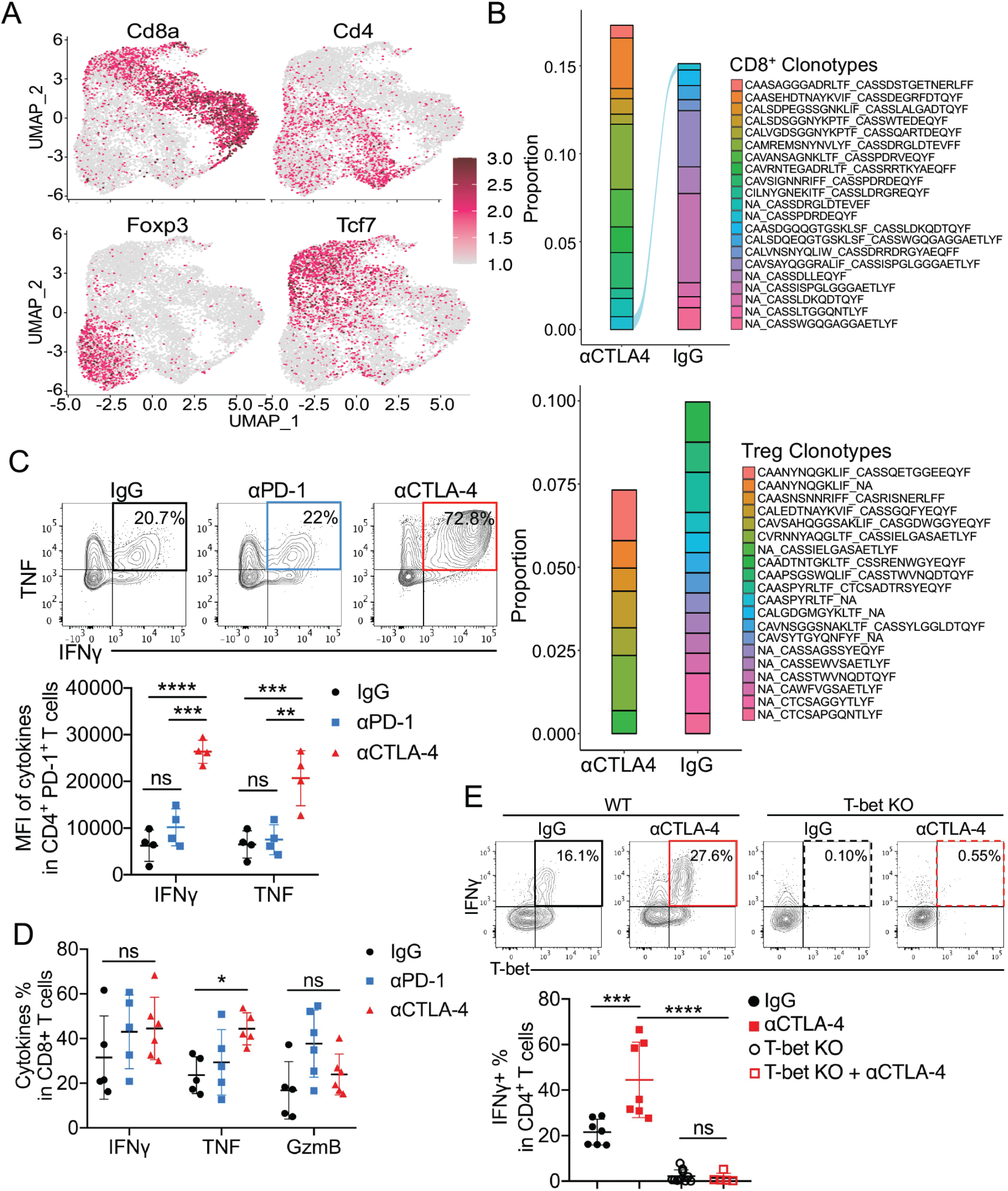
αCTLA-4 treatment increased the function of ‘Helper’ CD4^+^ T cells. (A) Feature plot showing the identical markers for general cell types. (B) Shared clonotype analysis in CD8^+^ T cells (above) or Treg (below) between IgG vs. αCTLA- 4 treatment by alluvial clonotypes. (C-D) C57BL/6 mice were implanted with 3 × 10^5^ 005 GSCs on day 0, treated with 10 mg/kg αPD-1, αCTLA-4 or IgG injected i.p. on days 7, 14, 21, 28, and euthanized on day 35. The expression of IFNγ^+^TNFα^+^ in CD4^+^ T cells from harvested brains were measured by flow cytometry (above), and the mean fluorescence intensity (MFI) of IFNγ^+^ and TNFα^+^ in CD4^+^ T cells from different treatment groups in 005 tumors were quantified by flow cytometry (below) (C). The expression of IFNγ^+^, TNFα^+^, GzmB^+^ in CD8^+^ T cells from harvested brains were measured and quantified by flow cytometry (D). n = 5 mice / group. (E) The expression of IFNγ and T-bet was measured in T-bet^+/+^ or T-bet^-/-^ CD4^+^ T cells from 005 tumors (above). Frequency of T-bet^+^IFNγ^+^ CD4^+^ T cells in the indicated groups was quantified (below).

**Figure S3.**
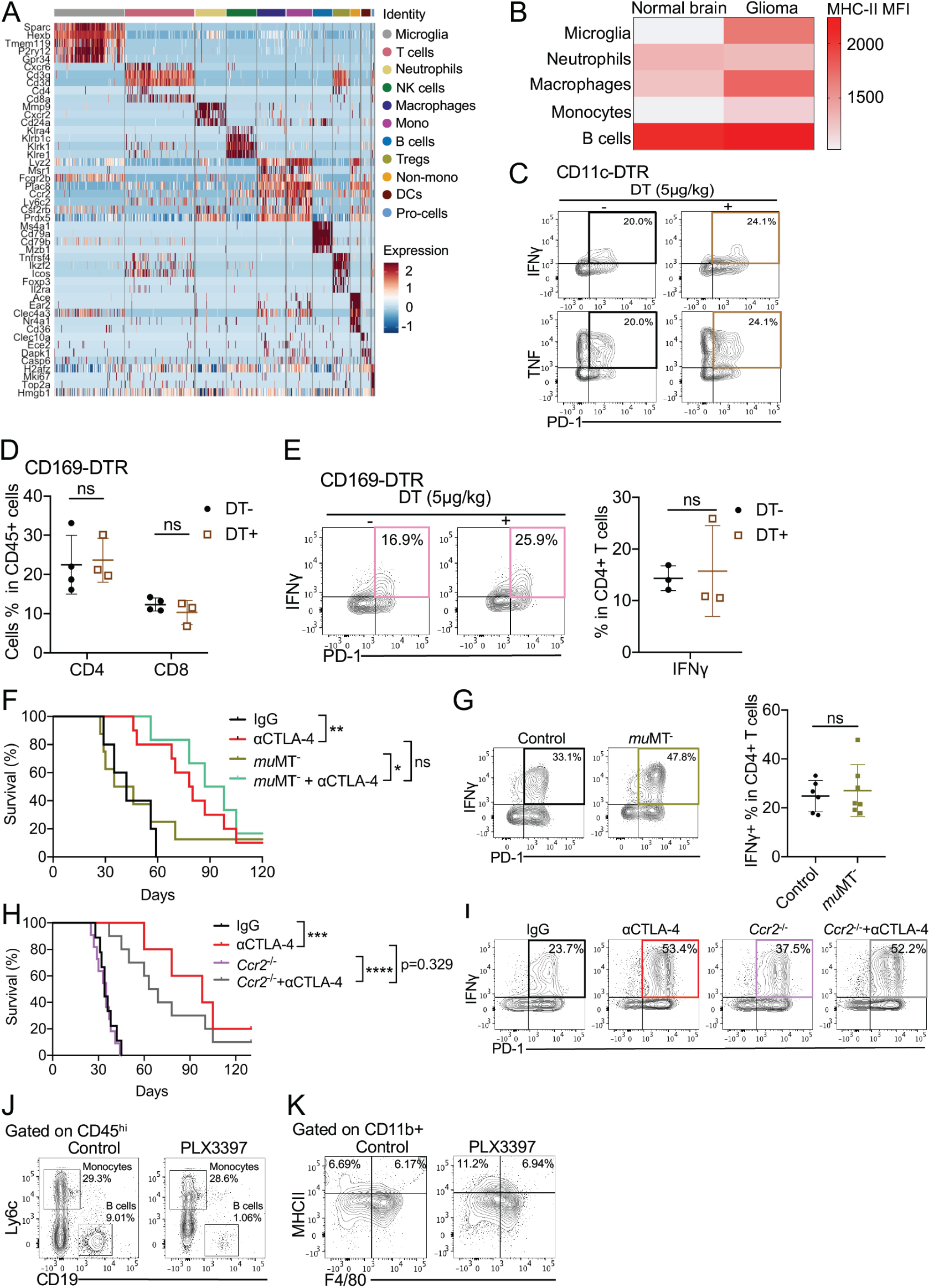
Anti-glioblastoma immune response does not depend on B cells or monocytes/macrophages. (A) Heatmap showing normalized expression of selected genes in each cluster. (B) The MFI of MHC-II in different subsets from normal brain or glioma were measured by flow cytometry. (C) Representative flow plots of the frequency of PD-1^+^IFNγ^+^ and PD-1^+^TNFα^+^ in CD4^+^ T cells from glioma tissues of CD11c-DTR mice with or without DT treatment. (D-E) CD169-DTR mice were administered with 5 μg/kg DT (i.p.) 2 weeks after glioma implantation, and maintained the treatment twice/week for 2 weeks. The percentages of CD4^+^ and CD8^+^ T cells (D), and PD-1^+^IFNγ^+^ of CD4^+^ cells (E), in brain tumor tissues from different treatment groups were measured and quantified by flow cytometry. (F) C57BL/6 or *mu*MT^-^ mice were implanted with 3 × 10^5^ 005 GSCs on day 0, treated with 10 mg/kg IgG or αCTLA-4 on days 7, 14, 21 and 28, and monitored for survival. Median survival of IgG (42 days; n = 5) was compared with αCTLA-4 (79 days; n = 10, p = 0.00114). Median survival of *mu*MT^-^ (40.5 days, n=9) was compared with *mu*MT^-^ treated with αCTLA-4 (92.5 days; n = 8, p = 0.0267). Similarly, αCTLA-4 treatment of wild-type B6 mice was compared with *mu*MT^-^ treated with αCTLA-4 (p = 0.4370) by log-rank analysis. (G) The expression of PD-1^+^IFNγ^+^ was measured in CD4^+^ T cells in control or *mu*MT^-^ mice from 005 tumors (left). Frequency of PD-1^+^IFNγ^+^ from CD4^+^ T cells in the indicated groups were quantified (right). (H) C57BL/6 or *Ccr2^-/-^* mice were implanted with 3 × 10^5^ 005 GSCs on day 0, treated with 10 mg/kg IgG or αCTLA-4 on days 7, 14, 21 and 28, and monitored for survival. Median survival of IgG (34 days; n = 9) was compared with αCTLA-4 (98 days; n = 5, p = 0.00065). Median survival of *Ccr2^-/-^* (35 days, n=11) was compared with *Ccr2^-/-^* treated with αCTLA-4 (66 days; n = 10, p = 0.000016). Similarly, wild-type B6 mice treated with αCTLA-4 were compared with *Ccr2^-/-^* treated with αCTLA-4 (p = 0.3292) by log-rank analysis. (I) The frequency of PD-1^+^IFNγ^+^ from CD4^+^ T cells in the indicated groups were measured by flow cytometry. (J) The frequency of monocytes (CD11b^+^Ly6c^+^) and B cells (CD19^+^) in control or PLX3397 treated groups were measured by flow cytometry. (K) The frequency of MHC-II^+^ in CD11b^+^ population from control or PLX3397 treated groups were measured by flow cytometry.

**Figure S4.**
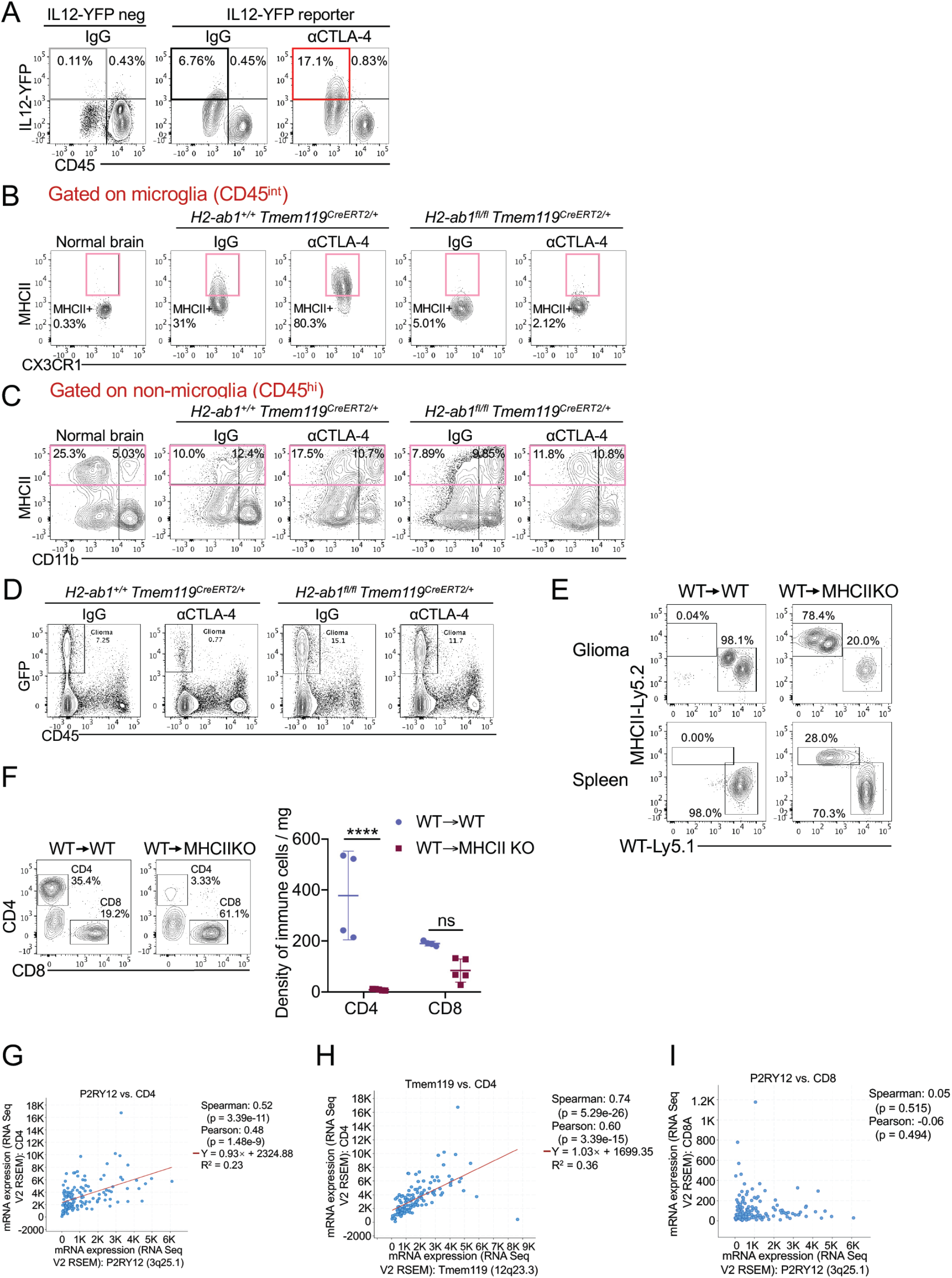
Microglia are required for CD4^+^ T cells activation in glioma. (A) IL-12^YFP^ reporter mice were implanted with 3 × 10^5^ 005 GSCs on day 0, treated with 10 mg/kg αCTLA-4 or IgG injected i.p. on days 7, 14, 21, 28, and euthanized on day 35. The representative flow plots of IL-12 were measured by flow cytometry in CD45^+^ cells from harvested brains. (B-D) *H2-ab1^fl/fl^Tmem119^CreERT2/+^* mice were treated with 200 mg/kg tamoxifen for 1 week, implanted with 3 × 10^5^ 005 GSCs on day 0, and treated with 10 mg/kg IgG or αCTLA-4. Brain tissues were harvested. The representative flow plots of MHC-II expression on microglia (CD45^int^) (B), non-microglia population (CD45^hi^) (C), and the density of glioma cells (CD45^-^ GFP^+^) (D) were checked by flow cytometry. (E-F) Bone marrow (BM) chimeras were generated by using C57BL/6 (Ly5.1) BM injected into Ly5.1 or MHC-II KO (C57BL/6 genetic background) mice at 5 × 10^6^ cells per recipient 4 hours after lead shield brain-protected 1200 Rads irradiation. Donor (Ly5.1^+^)-derived cells were reconstituted ten weeks after bone marrow transplantation (BMT) and the reconstitution efficacy was check by flow cytometry (E). Brain tissues were harvested on day 35 after 005 implantation, and the density of CD4^+^ and CD8^+^ T cells were measured by flow cytometry (F). (G-I) Dot-plots from TCGA data for glioblastoma to illustrate correlations between CD4 and P2RY12 (G), CD4 and Tmem119 (H), CD8 and P2RY12 (I).

**Figure S5.**
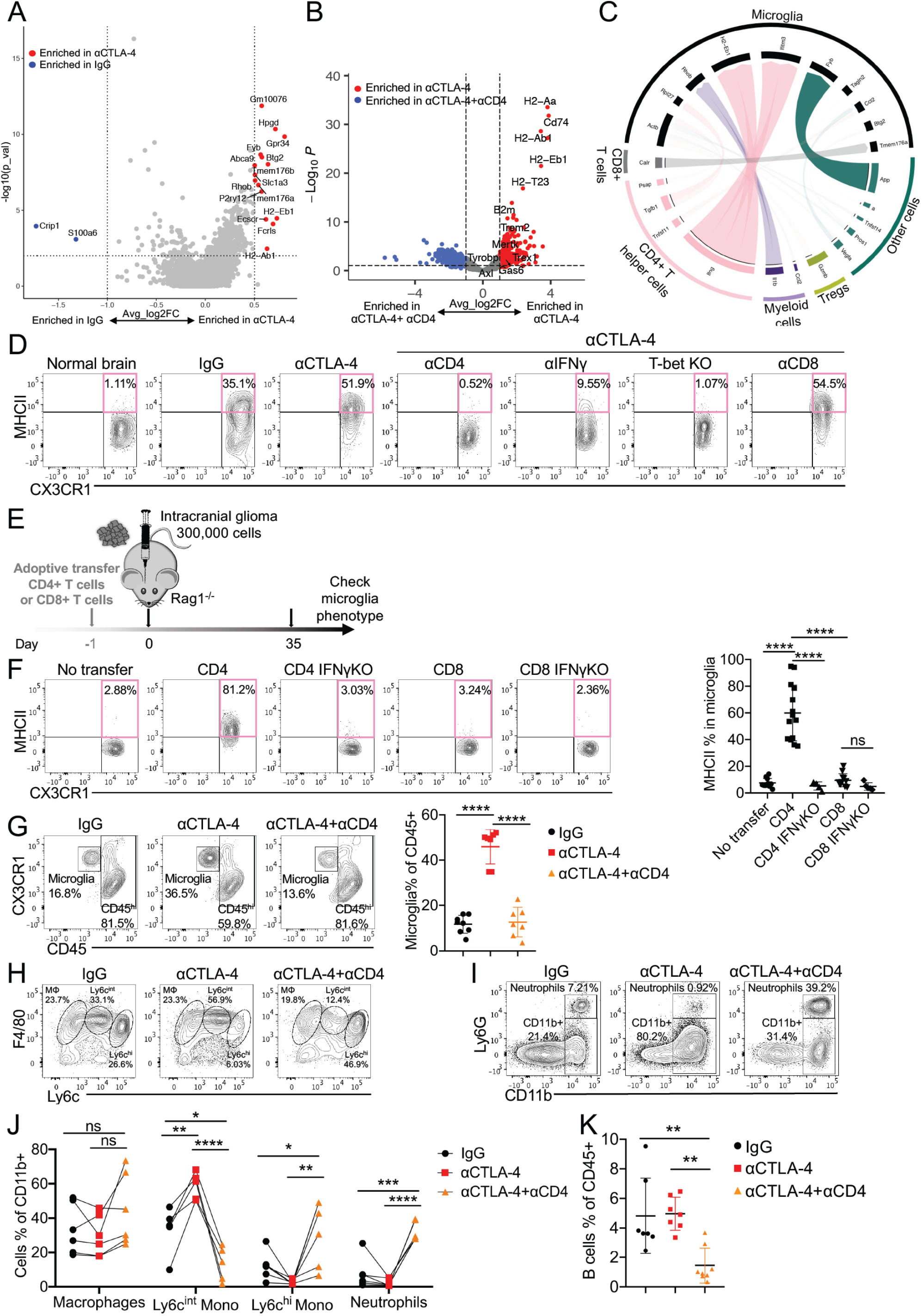
CD4^+^ T cells reprograming microglia and myeloid subsets. (A) Volcano plots of genes present the magnitude (log2 (fold change), x-axis) and significance (− log10 (adjusted P value), y-axis) for microglia from IgG versus αCTLA-4 from sc-RNAseq. Each spot represents a transcript. Two vertical dashed lines represent the threshold of fold changes (log2 (fold change)>0.5 or< −1). (B) Microglia (CD45^int^CX3CR1^+^) were sorted from 005 glioma-bearing brains treated with IgG, anti-CTLA-4, or αCTLA4 combined with CD4 depletion and then subjected to bulk RNA-seq. Volcano plots of genes present the magnitude (log2 (fold change), x-axis) and significance (− log10 (adjusted P value), y-axis) for microglia from αCTLA-4 vs. αCTLA-4+αCD4 from bulk RNA-seq. (C) NicheNet circle plot showing links between upregulated cytokines and chemokines in CD4+ (pink), CD8+ (gray), myeloid cells (purple), Tregs (green) and other cells (teal), with upregulated genes in microglia (black) after αCTLA-4 treatment. (D) Representative flow plots of Figure 5F. (E-F) Purified naïve CD8^+^CD45.1^+^ or CD4^+^CD45.1^+^ cells were electroporated with targeting or non-targeting *Ifng* sgRNA/Cas9 RNPs and 1x10^6^ cells were immediately transferred into Rag1^-/-^ recipient mice on day 0. All the recipient mice were implanted with 3 × 10^5^ 005 GSCs on day 1 (E). MHC-II expression on microglia within tumor tissue were analyzed and quantified on day 28 (F). (G-K) C57BL/6 mice were implanted with 3 × 10^5^ 005 GSCs on day 0, and treated with 10 mg/kg isotype IgG or αCTLA-4 with or without αCD4 antibody injected i.p. on days 7, 14, 21 and 28. The percentages of CD45^int^CX3CR1^+^ microglia cells in different treatment groups from 005 tumors were measured and quantified by flow cytometry (G). Representative flow plots of the frequency of macrophages (F4/80^+^), monocytes (Ly6c^hi^), Ly6c^int^ in CD11b^+^Ly6G^-^ cells (H); neutrophils (% of CD11b^+^Ly6G^+^ cells in CD45^+^ cells) (I). The corresponding quantification summary from H-I is shown (J). Representative quantification (K) of B cell frequencies from tumor tissues in different treatment groups. All results were pooled from or representative of 2-4 experiments with n = 3-5 mice / group (A), n = 3 mice / group (B), n = 3 mice / group (C), n = 5-8 mice / group (D), n = 4-12 mice / group (F), n = 6-8 mice / group (G-K). Data are expressed as mean ± SEM. Statistical analysis was performed using Student’s *t* test (two tailed) comparing IgG group to various treatment groups. * p < 0.05, ** p < 0.01, *** p < 0.001, **** p < 0.0001.

**Figure S6.**
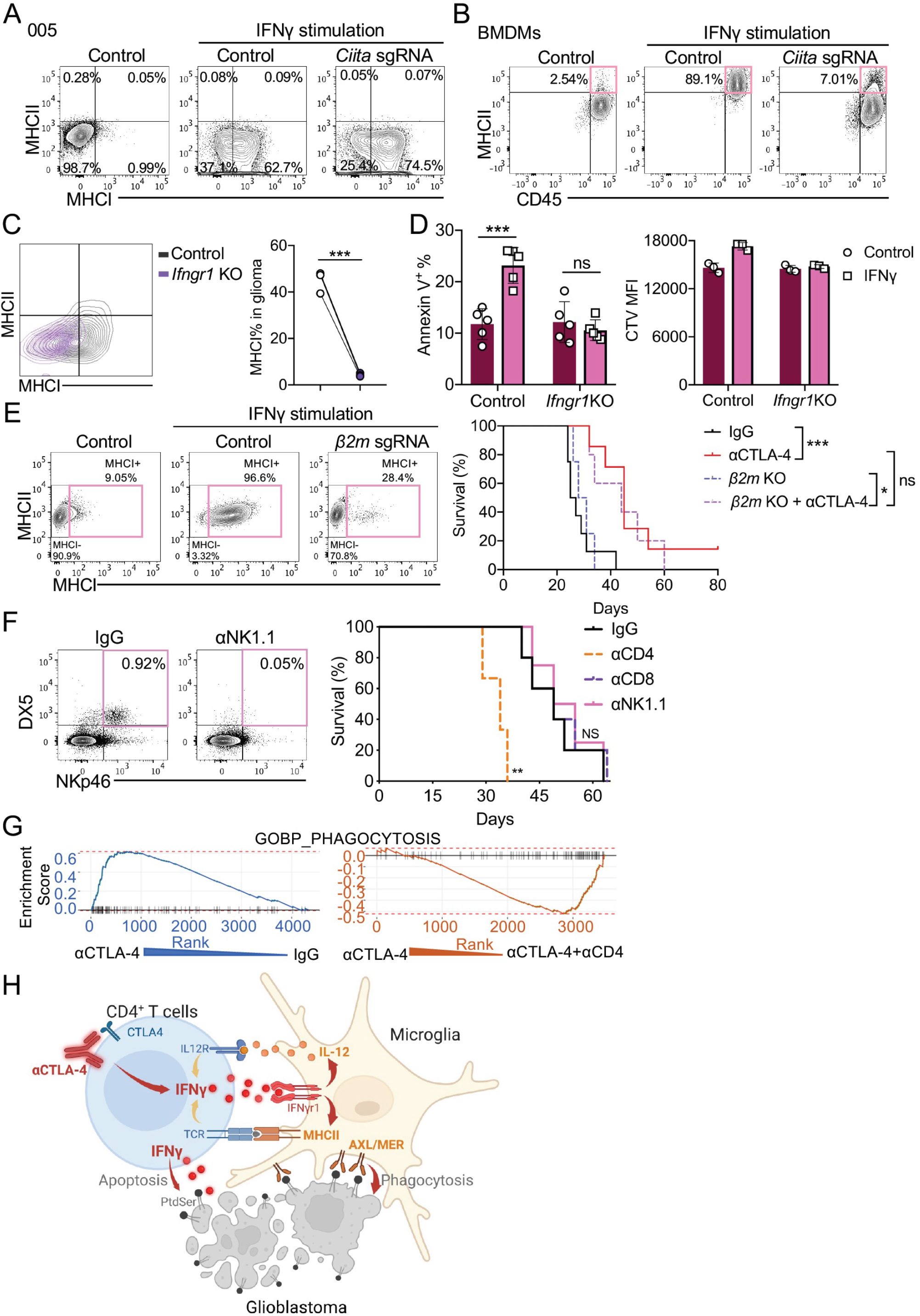
CD8^+^ T cells and NK cells were not involved in anti-glioma immunity, but CD4^+^ T cells regulated phagocytosis genes on microglia. (A) 005 cells were electroporated with targeting or non-targeting *Ciita* sgRNA/Cas9 RNPs to generate *Ciita* KO (MHC-II KO) 005 cell line. (B) Deletion efficiency of same targeting or non-targeting *Ciita* sgRNA/Cas9 RNPs (A) was validated in the high MHC-II expressing bone marrow derived macrophages (BMDMs). (C) 005 cells were electroporated with targeting or non-targeting *Ifngr1* sgRNA/Cas9 RNPs to generate *Ifngr1* KO 005 GSCs. C57BL/6 mice were implanted with 3 × 10^5^ WT or *Ifngr1* KO 005 GSCs and euthanized on day 35. MHC-II and MHC-I expression on glioma were analyzed and quantified by flow cytometry. (D) WT or *Ifngr1* KO 005 cells were treated with IFNγ (10 ng/ml) for 24 hours. The apoptosis cells (Annexin V^+^) and proliferation (CTV) were measured and quantified by flow cytometry. (E) 005 cells were electroporated with targeting or non-targeting *B2m* sgRNA/Cas9 RNPs to generate β*2m* KO (MHC-I KO) 005 cell line. Mice were implanted with 3 × 10^5^ 005 or β*2m* KO 005 on day 0, treated with 10 mg/kg IgG or αCTLA-4 injected i.p. on days 7, 14, 21 and 28, and monitored for survival. Median survival of IgG (26 days; n = 8) was compared with αCTLA-4 (45 days; n = 7, p = 0.0007). Median survival of β*2m* KO (29.5 days, n=4) was compared with β*2m* KO treated with αCTLA-4 (44 days; n = 5, p = 0.024). Similarly, comparison of αCTLA-4 treatment of WT 005 with that of β*2m* KO 005 (p = 0.5645) were done by log-rank analysis. (F) Mice were implanted with 3 × 10^5^ 005 GSCs on day 0, and treated with 10 mg/kg IgG, αCD4, αCD8, αNK1.1 antibodies, injected i.p. on days 7, 14, 21 and 28. Median survival of IgG (49 days; n = 5) was compared with αCD4 (34 days; n = 5, p = 0.004), αCD8 (49 days; n = 5, p < 0.6370), αNK1.1 (52 days; n = 4, p = 0.6233) by log-rank analysis (right). Representative flow plots of NK cells (DX5^+^CD335^+^) depletion in brain (left). (G) The analysis of gene set enrichment analysis (GSEA) demonstrated known phagocytosis pathway, were enriched in αCTLA-4 compare to IgG (blue). Negative regulation of phagocytosis pathway are enriched in αCTLA-4+ αCD4 compare to αCTLA-4 (red). (H) Scheme explaining the partnership between microglia and CD4^+^ T cells. All results were pooled from or representative of 2-4 experiments with n = 3 mice / group (C). Data are expressed as mean ± SEM. Statistical analysis was performed using Student’s *t* test (two tailed) comparing IgG group to various treatment groups. * p < 0.05, ** p < 0.01, *** p < 0.001, **** p < 0.0001.

